# Long-read sequencing of *Mycobacterial tuberculosis* is comparable to short-read sequencing for antimicrobial resistance prediction and epidemiological studies

**DOI:** 10.64898/2026.04.08.717216

**Authors:** Matthew Colpus, Catriona S Baker, Eloïse Roghi, Hanh Nguyen Hong, Phu Phan Trieu, Do Dang Anh Thu, Alexandra Hall, Philip W Fowler, Timothy M Walker, Ruan Spies, Hermione Webster, Jeremy Westhead, Hieu Thai, Robert D Turner, Timothy EA Peto, Nguyen Le Quang, Nguyen Thuy Thuong Thuong, Shaheed Valley Omar, Derrick W Crook

## Abstract

**Background:** Short-read genetic sequencing technologies (mainly Illumina) have been extensively used for around a decade for *Mycobacterium tuberculosis* complex (MTBC) outbreak analysis and genomic drug susceptibility testing (gDST) with the result that Illumina has become the *de facto* gold standard. Long-read sequencing, as exemplified by Oxford Nanopore Technologies (ONT), offer the prospect of faster, simpler, and portable sequencing. In this work, we carry out the largest to date comparison of how well Illumina and ONT technologies sequence MTBC samples, making use of R10.4 ONT flowcells, updated basecalling models and deep-learning variant calling.

**Methods:** A total of 508 samples were sequenced using both short and long-read platforms. All samples originated from South Africa or Vietnam and were over-selected for drug resistance and also included several local outbreaks and a range of lineages. The South African and Vietnamese samples had already been Illumina sequenced. Samples with ≥50 read depth by Illumina were selected for sequencing by ONT using one of the GridION or PromethION platforms. Bioinformatics processing was done using a modified online cloud platform which included reference-based variant calling, catalogue-based gDST and identified related samples via SNP counting to inform outbreak detection. The lineages and gDST predictions obtained by short-and long-sequencing were compared for all samples as were all putative clusters identified via SNP counting. For convenience Illumina was used as the reference method.

**Findings:** Of the 508 samples, 425 (83.7%) had sufficient read depths to permit comparison between the two sequencing technologies. The assigned lineages were identical for 407/425 (95.8%) samples and all discordances were due to mixed lineages being identified by one technology. Evidence of non-tuberculous mycobacterium (NTM) subpopulations were found in nine samples. Using Illumina as the reference method, the very major error (VME) rate of ONT for predicting resistance to all 15 drugs is 1.0% (0.6-1.5%) whilst the major error (ME) rate is 1.7% (1.3-2.2%) with an unclassified rate of 6.9% (6.3-7.5%). This is below the thresholds specified by the CLSI. Considering each of the 15 drugs individually they had VME and ME point estimates below ≤3% in 29/30 cases; and most 25/30 below ≤1.5%. Filtering out all samples containing mixtures left 382 isolates. By appropriate masking of the reference genome we were able to obtain a mean SNP distance between the two platforms of 0.13 (median of zero) for the same sample and for 376/382 samples (98.4%, CI:96.6-99.4%) the difference was ≤1 SNPs. The high concordance in SNP identification ensured that few differences in the 43 putative clusters among 172 isolates were observed.

**Interpretation:** The differences between the two sequencing platforms for the key clinical outputs is so small that it is now within the tolerances set by regulatory agencies. Provided the sequencing is of sufficient quality, we have therefore reached a threshold whereby sequencing data from long-and short-read platforms can be aggregated. This will enable large scale analyses by national and international public health agencies whilst allowing the MTBC community to take advantage of the portability and speed of long-read sequencing.

**Funding:** The NIHR Health Protection Research Unit: Healthcare Associated Infections and Antimicrobial Resistance at University of Oxford (NIHR200915), a partnership between the UK Health Security Agency (UKHSA) and the University of Oxford, the National Institute for Health and Care Research Biomedical Research Centre: Oxford (BRC) and the Ellison Institute of Technology, Oxford Ltd. The CRyPTIC project was funded by Wellcome [214560/Z/18/Z], a Wellcome Trust/Newton Fund-MRC Collaborative Award (200205/Z/15/Z); and the Bill & Melinda Gates Foundation Trust (OPP1133541).

**Research in context:** *Evidence before this study:* We conducted a PubMed Central full text search for “tuberculosis” AND (“drug resistance prediction” OR “drug susceptibility prediction”) AND (“genome” OR “genomic” OR “genotypic”) AND (“ont” OR “oxford nanopore”) between 2022 and 2026 (conducted 1 April 2026). This returned 62 papers; of which, six used both Illumina and ONT sequencing. One of these, published in 2023, directly compared the performance of the two platforms on 151 M. tuberculosis isolates oversampled for resistance. The investigation yielded comparative results for the earlier generation ONT flow cell (R9·4·1) and base-caller (guppy version 5·0·16). Another, published in 2026, investigated a targeted next-generation sequencing panel of 20 amplicons using ONT sequencing on R10.4.1 flow cells with guppy 6·4·6. They compared the results on 71 isolates against phenotypic data and Illumina whole genome sequencing (for 53 isolates) but had low rates of resistance, with all drugs but isoniazid being limited to under five resistant isolates. Two other small studies (10 and 13 samples, respectively) conducted feasibility studies comparing ONT with Illumina, also using earlier generation flow cells and base-calling technology from ONT. Two further studies compared Illumina with ONT for direct sputum sequencing and did not investigate the comparative performance of the two platforms for variant call accuracy, resistance prediction, and outbreak detection. Illumina sequencing technology is widely used for genomic sequence analysis in research, and clinical and public health contexts. Consequently, it has become the *de facto* reference standard for generating whole genome sequence data. Whilst previous studies established the promise and limitations of long-read (ONT) sequencing as an alternative to short-read sequencing (mainly Illumina), the enhanced performance arising from newer flowcells (e.g. R10.4.1), V14 chemistry, and the latest basecallers (dorado v4.3.0/5.0.0) has not been analysed. Neither has any ONT analysis incorporating the new deep-learning variant callers been evaluated in a large-scale comparative study. Thus, it is currently unclear whether data generated by either platform can be used safely in aggregated analyses for research and clinical or public health service.

*Added value of this study:* We compared how well short-(Illumina) and long-read (ONT) sequencing platforms identify the genetic variants in *M. tuberculosis*, predict antituberculous drug resistance and recog-nise outbreaks. The long-reads were generated using the latest generation ONT R10.4.1 flows cells, V14 chemistry, ‘super high accuracy’ basecalling (dorado v4.3.0/5.0.0) and a bioinformatics analysis pipeline built using the Clair3 deep-learning based variant caller. A total of 508 clinical samples were sequenced using both technologies, substantially more than previous studies. The sampling frame was much larger than previously investigations and included a large proportion of isolates with resistance to first-line and second-line antibiotics as well as bedaquiline. Thus, providing greater statistical power for resistance prediction than before. In particular, the inclusion of bedaquiline resistance provided evidence useful for predicting resistance to this newly deployed drug for treating multi-drug resistant (MDR) TB. We find that the differences between technologies are small meaning that either technology can be used alone safely, and services using both technologies can confidently aggregate the data for analysis.

*Implications of all the available evidence:* This will be a benefit to local, regional and international organisations, particularly public health agencies, which often have a mix of the two main sequencing technologies for characterising TB whole genome sequences. It also opens up the sequence based diagnostic market to greater competition, particularly if the observed performance can be replicated for other pathogen species.

## Introduction

Over 10 million cases of tuberculosis (TB) were reported in 2024 and 1.23 million people died [1] confirming the aetiological agent, *Mycobacterium tuberculosis*, as the single most deadly pathogen on our planet. The WHO recommends using whole genome sequencing (WGS) or targeted next generation sequencing (tNGS) for surveillance of drug-resistant TB [2] and tNGS for detecting drug resistance in individuals [3].

To date the majority of TB genetic research and surveillance has used short-read sequencing platforms (such as those produced by Illumina, Inc.) due to their low error rate, enabling single-nucleotide polymorphisms (SNPs) and small insertions or deletions (indels) to be reliably detected. For example, as of March 2026, the vast majority of sequencing runs (314,619 / 324,347, 97.0%) deposited in the European Nucleotide Archive that purport to contain *M. tuberculosis* were sequenced using an Illumina platform. Illumina sequencing can therefore be considered as the *de facto* reference standard against which other sequencing technologies should be assessed.

Long-read sequencing platforms, as exemplified by those manufactured by Oxford Nanopore Technologies plc (ONT) or Pacific Biosciences of California, Inc (PacBio), offer the additional benefit by being able to better resolve repetitive regions (e.g. the PE/PPE gene family) or detect large structural variants [4]. However, the adoption of long-read sequencing platforms for clinical or public health use has been inhibited by the paucity of studies demonstrating near equivalent performance to short read platforms. ONT is also potentially more portable than the other platforms which would be of great benefit to healthcare or public health units in less accessible areas, potentially extending the reach of sequencing technologies for TB management to those places where it can have the greatest health benefits.

In 2023 Hall *et al.* [5] compared 151 samples sequenced using both Illumina and ONT plat-forms, and concluded that ONT sequencing showed great promise. Despite only being three years ago, there have since been advancements in flowcells, basecalling software and variant calling; they used R9.4.1 flowcells, guppy version 5·0·16 and BCFTools [6], respectively. It is therefore timely to undertake a comparative study to assess whether ONT sequencing has matured to the point such that it can be used as a reliable alternative technology to Illumina. In this study, we will evaluate the performance of long-read (ONT) sequencing at (1) identifying mutations associated with antimicrobial resistance and (2) assessing if two samples are sufficiently similar that they are likely part of the same outbreak as was previously identified via short-read (Illumina) sequencing.

## Methods

### Sample collection and sequencing

A total of 508 clinical sputum samples were collected between March 2020 and April 2024 from South Africa (n=317, with three H37Rv controls) and Vietnam (n=191, with one H37Rv control). After standard decontamination procedures all samples were cultured in Mycobacterial Growth Indicator Tubes (MGIT; Becton Dickinson [BD], Franklin Lakes, NJ, USA) using the BACTEC MGIT 960 system (BD, Franklin Lakes, NJ, USA) for automated growth detection. After flagging positive, cultures were sub-cultured onto Löwenstein–Jensen (LJ) agar slants (BD, Franklin Lakes, NJ, USA) to obtain mature colonies. DNA from the South African isolates was extracted [7] and Illumina sequenced [8] as previously described. The Vietnamese isolates were extracted using the cetyl trimethylammonium bromide method [9].

Samples from both countries were selected for ONT sequencing if they had produced at least 50-fold read depth with Illumina. All South African isolates were sequenced using a modified SQK-RPB114.24 method as previously described [10] on a GridION using R10.4.1 flow cells with V14 chemistry. Samples from Vietnam were selected if, in addition, their DNA concentration was *>*20 ng*/*µL and were sequenced by a commercial supplier. DNA was quantified using a Qubit 4.0 Fluorometer and qualified using an Agilent 5600 Fragment Analyzer. ONT library preparation was performed with the Rapid Barcoding Kit 96 V14 according to the manufacturer’s recommendations. Sequencing was run on PromethION flow cells (R10 M Version) on a PromethION 2 Solo for 72 hours. Read filtering was conducted with a minimum quality score of Q9 and minimum read length of 1kb. Due to read depth issues, 81 Vietnamese samples were repeated in-house using the same protocol as the South African samples. All basecalling was performed with dorado using either v4.3.0 or v5.0.0 (Table S1).

### Bioinformatics processing

Sequencing data was processed using an in-house variant of an online cloud platform [11]. All samples had human reads removed using Hostile (v1.1.0) [12] or Deacon (v0.7.0) [13], before filtering reads based on quality with Fastp (v0.24.1) [14]. Kraken2 (v2.1.3) was used to select reads from the *Mycobacteriaceae* family (using the 6/5/2023 Standard index) [15]. The reads were then mapped using Minimap2 (v2.28) against a multifasta of *Mycobacterium* reference genomes to select the TB reads and determine contaminating non-tuberculous mycobacterial species, whilst Mykrobe (v0.13.0) was used to determine the precise lineage [16, 17].

Variants with respect to the H37Rv (v3) reference were identified using Clockwork (v0.12.4) for Illumina [18, 19], and Clair3 (v1.1.2) [20] for ONT, with BCFTools (v1.22) [21] used to determine read depth, at sites not reported by Clair3, for the purpose of reporting null calls. Gnomonicus (v3.1.1) was used to predict resistance with version 2 of the WHO catalogue (WHOv2) [11, 22]. Variants were included for resistance prediction provided they had ≥5% of the read support at that site and ≥5 reads for ONT or ≥3 reads for Illumina. Those with ≥90% of the read support were considered ‘major’ variants, while the rest were labelled ‘minor’ variants. Only ‘major’ variants were added to the consensus fasta file, which was then used in the relatedness analysis. Any genetic loci with ≥2 variants (above the minimum thresholds just specified) was counted as ‘mixed’ for the purpose of detecting mixed strains. Uncatalogued mutations in resistance determining genes were classified as ‘Unknown’, while ‘Fail’ was returned if there was insufficient depth at a site or it was otherwise filtered (Supplement §5). To guide interpretation we treated the short-read sequencing data as the gDST reference method to which we compared the ONT data. This allows us to apply the CLSI standard [23] which states that a DST method should have a very major error rate (false negative rate) ≤ 1.5 % and a major error rate (false positive rate) ≤ 3 % relative to the reference standard.

Whether two samples were epidemiologically related was determined by the magnitude of the SNP distance between them. A mask was first applied to both genomes to filter out repetitive and low complexity regions, as well as regions that result in frequent errors when mapping Illumina reads [24]. We assessed the mask by looking at positions with differences between the two platforms in multiple samples, and consequently, added nine extra positions to the mask (Supplement §8.1). Only SNPs without another SNP within 12 bases were included in the SNP distance calculation (Supplement §8.3 for an evaluation of this filter) [25].

Since each sample can be sequenced with either ONT or Illumina, there are four permutations and hence four distances for each pair of samples. We define the *shared SNPs* as the number of positions with a SNP between the two samples in all permutations. The Clopper-Pearson exact binomial method was used to calculate 95% confidence intervals.

### Sample filtering

Sample were excluded due to failing to PCR in the modified SQK-RPB114.24 sample preparation (n=6), insufficient ONT reads (n=75), or sample mislabelling (n=2), leaving 425 samples for comparison (Fig. S1).

## Results

Of the 508 isolates, we successfully obtained long genetic reads from 425 samples with both ≥10X depth and ≥90% genome coverage. All samples had previously been short read sequenced (using an Illumina platform) to at least 50X depth. Three out of the 19 ONT flow cells had substantially lower yields (*<*100 Mbp per sample) and we therefore resequenced the 81 samples most affected. The final samples had a median sequencing yield of 643 Mbp with Illumina and 525 Mbp with ONT (Fig. S3).

### Identification of mixed infections

All samples were expected to contain MTBC, however we also found evidence of a nontuberculous mycobacterial (NTM) infection in nine samples (seven *M. avium*, one *M. novom* and one *M. marseillense*). In all cases, strong support was found by mapping ONT reads, whereas, with Illumina there was strong support in five and weak support in the remaining four samples (Supplement §4).

In 406 (95.5%) of the 425 samples the identified MTBC lineage and sub-lineages were concordant between the sequencing platforms. The 19 discordant samples were determined as containing a mixed TB infection by one or other of the technologies (Illumina n=17, ONT n=2). No sample was identified as a mixed TB infection on both platforms by this method; however, by counting genetic loci with evidence of heterogeneity (Supplement §6) we identified 43 putative mixed TB infections: 21 in both technologies (including the 17 identified by Illumina), four in Illumina alone and 18 in ONT only (including both ONT mixed-lineage samples). Notably all ONT-only mixed-strain samples came from the South African dataset. A total of 35 distinct sub-lineages were common to both platforms; these were spread across the main lineages (1-4) with 230/425 (54.1%) belonging to lineage 2.

### Detection of resistance-associated variants

We first considered all 536 genetic loci where a SNP could confer resistance with high confidence according to the WHOv2 catalogue (annotated ‘Group 1: Assoc w R’) [22]; this yields 227,800 comparisons across the 425 samples (Table S2, Supplement §7). As mentioned, treating the Illumina results as the reference standard allows us to calculate error rates as if ONT were a new gDST method [23]. ONT missed 35/1885 (false negative rate = 1.9%, CI:1.3-2.6%) variants detected by Illumina, of which 29 variants were minor SNPs and six variants were a result of a single deletion. ONT reported an additional 14 variants (false positive rate = 0.006%, CI:0.003-0.01%), 12 of which resulted from a single minor deletion leaving two further mutations.

The WHOv2 catalogue describes genetic variants as amino acid mutations, insertions, deletions or SNPs (for RNA-coding genes and promoter mutations) so we must now translate the resulting nucleotide sequences into amino acids, where appropriate. We find 2,736 mutations associated with resistance or which had an unknown classification (Table S3, Supplement §5). The majority, 95.6% (2,616/2,736), were identified by both platforms (SNPs=2,326, indels=290) of which 257 were only present as a minor variant in one or both platforms. That leaves 120 discrepancies; about half, 61 (2.2% of 2,736), were due to a deletion in a tandem repeat section of *fbiC* exclusively resolved by ONT (Supplement §7.1). Two indels were exclusive to Illumina: a 2 kb *katG* deletion (Supplement §7.2) and a two base pair deletion in *eis*. The remaining 57 were all minor SNPs (ONT=1, Illumina=27) or minor indels (ONT=2, Illumina=27) resolved by one platform but not the other. Short insertions or deletions in *Rv0678* accounted for 23 of the minor indels whereas minor SNPs were found in a number of genes. Again, treating Illumina as the reference method yields 64 false positives and 56 false negatives for ONT.

The apparent greater ability of Illumina to detect minor variants is likely explained by a combination of higher accuracy and/or greater read depths (which increases the chance of having enough reads carrying the minor variant for it to be detected). To investigate this we looked at the 27 minor SNPs resolved by Illumina only. In 26/27 cases there is evidence of the SNP in the ONT pileup but it is either below the arbitrary five read minimum threshold for calling (n=8) or is not called by Clair3 (n=18) despite there being sufficient reads.

The only differences resulting from major variants came from indels. Comparing across the whole genome, ONT finds evidence for 44960/46055 (97.6%) of the major indels reported by Illumina, whereas Illumina only finds evidence for 37867/45413 (83.4%) of those reported by ONT (Supplement §7.3, Tables S4, S5). This is consistent with the apparent greater ability of long-read sequencing to resolve structural variation.

### Antimicrobial resistance prediction

Next we compared the predicted resistant/susceptible phenotype for the 15 drugs included in the WHOv2 catalogue [22] (Fig. 2). Across all drugs a concordant result was obtained for 333/425 (78.4%) samples. The remaining 92/425 (21.6%) samples contained all the 127/6,375 (2.0%) discordant classifications. About half, 60/127 (47.2%), are due to a detected *fbiC* deletion triggering a resistance call in ONT for delamanid, though we suggest that the nature of the deletion would not in fact give rise to resistance (§7.1). Hence a small proportion of the samples, 38/425 (8.9%), give rise to the remaining 67/127 (52.8%) differences; 30 of these are due to a resistance-associated mutation being identified in one platform but not the other, five of these are due to the software reporting a fail due to insufficient reads at a genetic loci associated with resistance, and 32 due to a mutation with an unknown effect being reported by one platform.

**Figure 1:**
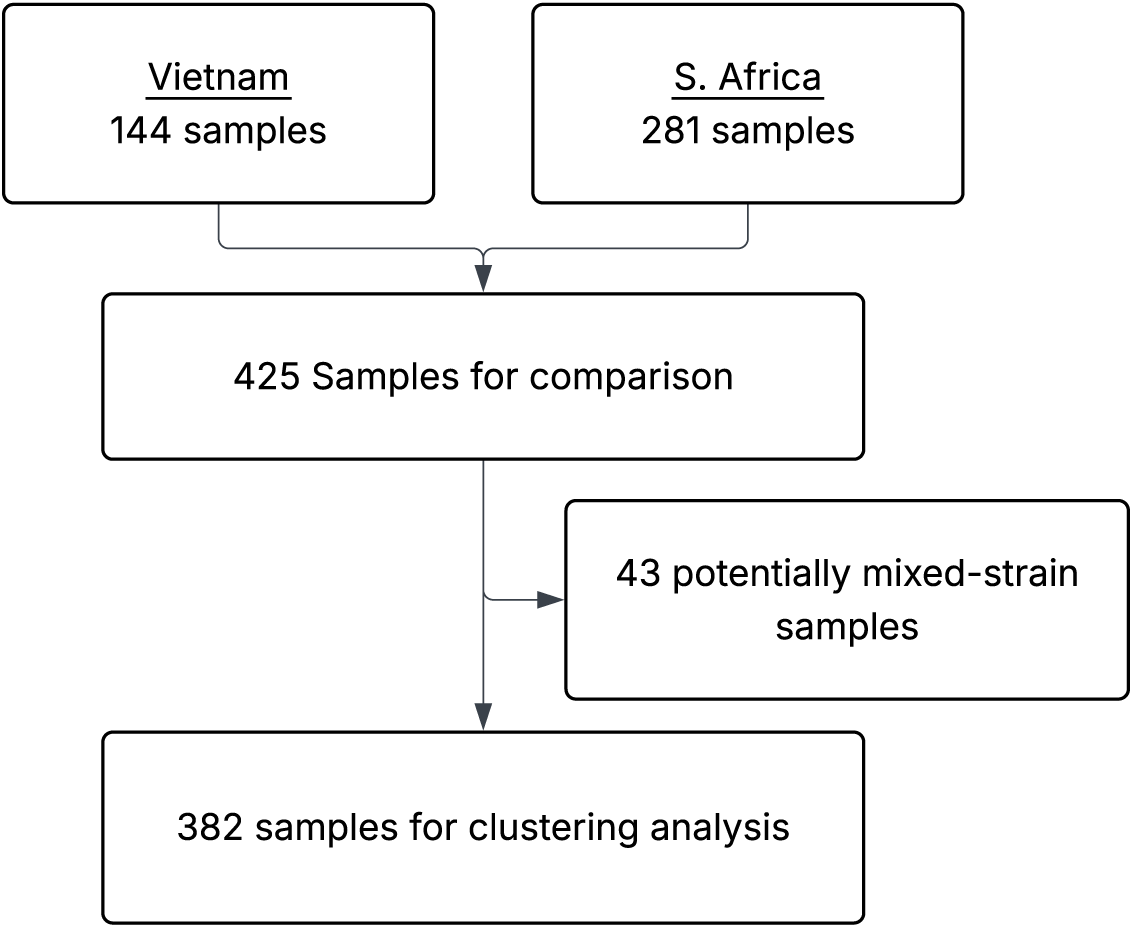
Sample collection and filtering.

**Figure 2:**
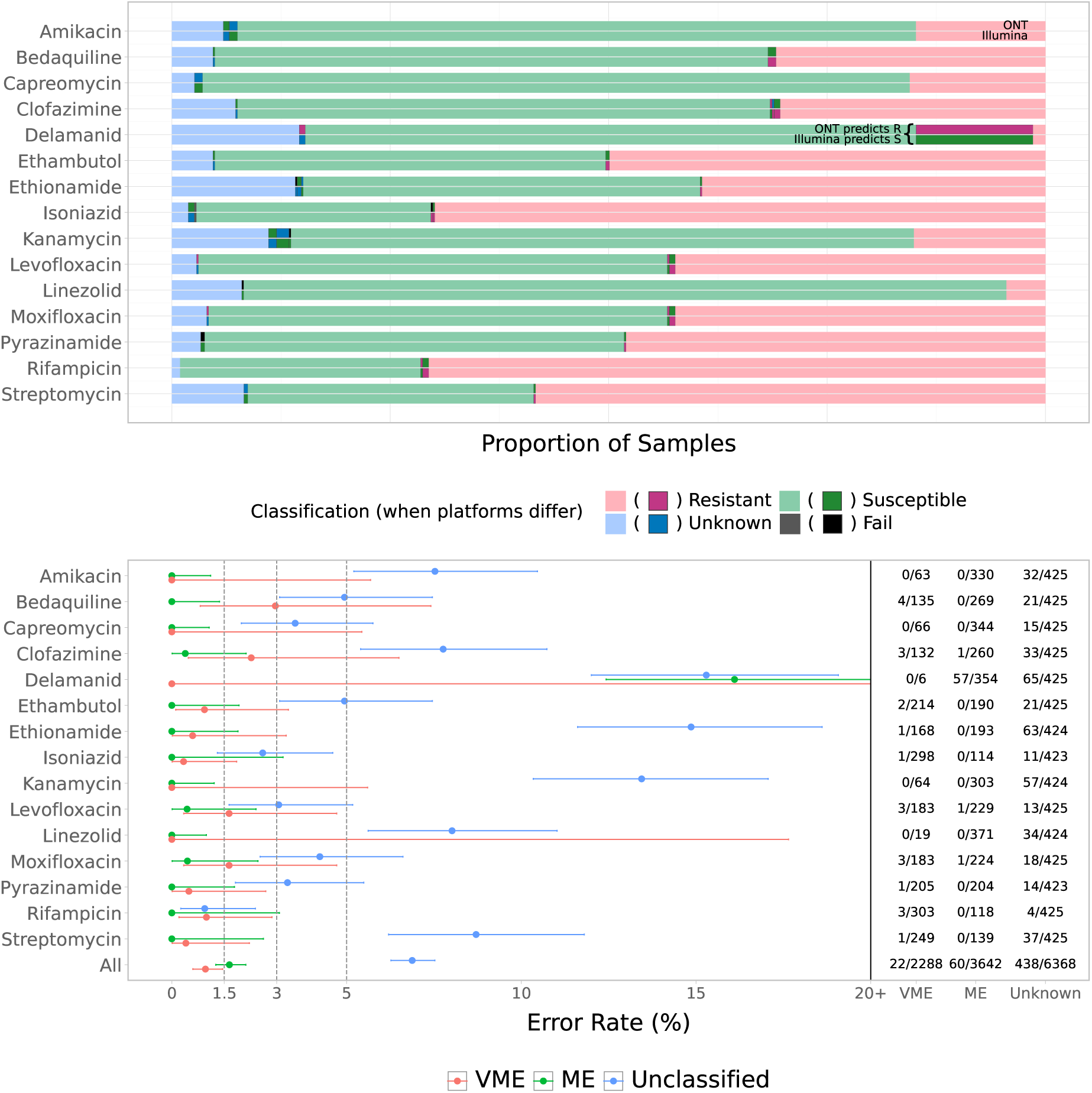
Comparison of genotypic resistance classification between ONT and Illumina. a) The major and very major error rates of ONT when using Illumina as the ‘true’ signal as well as the proportion of samples classified as unknown by either platform. b) For each drug the lower bar shows the proportions of samples classified as Unknown/Susceptible/Resistant by Illumina, and the top bar by ONT for the corresponding samples. Where platforms differ the sections are shown in bold with darker colours. i.e. The annotated part of delamanid shows that 14% of samples were considered susceptible by Illumina but resistant by ONT.

Treating Illumina sequencing as the reference method allows us to calculate the very major error rate (VME, false negative rate) and major error rate (ME, false positive rate) for ONT. Samples in which either platform assigns an unknown/failed classification are excluded from VME/ME calculations. Combining the classifications of all drugs across all samples yields an overall VME rate of 1.0% (0.6-1.5%) and an overall ME rate of 1.7% (1.3-2.2%), with an unclassified rate of 6.9% (6.3-7.5%). Despite there being variability between the different drugs, all had VME rates ≤3% with 11/15 being ≤1.5%. Of those 11, six have 95% confidence intervals below 5% and the other five had no very major errors but too few resistant samples to meet this bound. All drugs, except delamanid, had ME rates ≤1.5% with the 95% confidence interval below 5%.

### SNP differences between platforms

To investigate the SNP distances, we first excluded samples with evidence of being mixed strains, leaving 382 samples for all subsequent analysis. After masking (Supplement §8.1), the mean SNP distance between platforms on the same sample was 0.13 (median of zero), with 376/382 (98.4, CI:96.6-99.4%) being ≤1 SNPs (Fig. 3).

**Figure 3:**
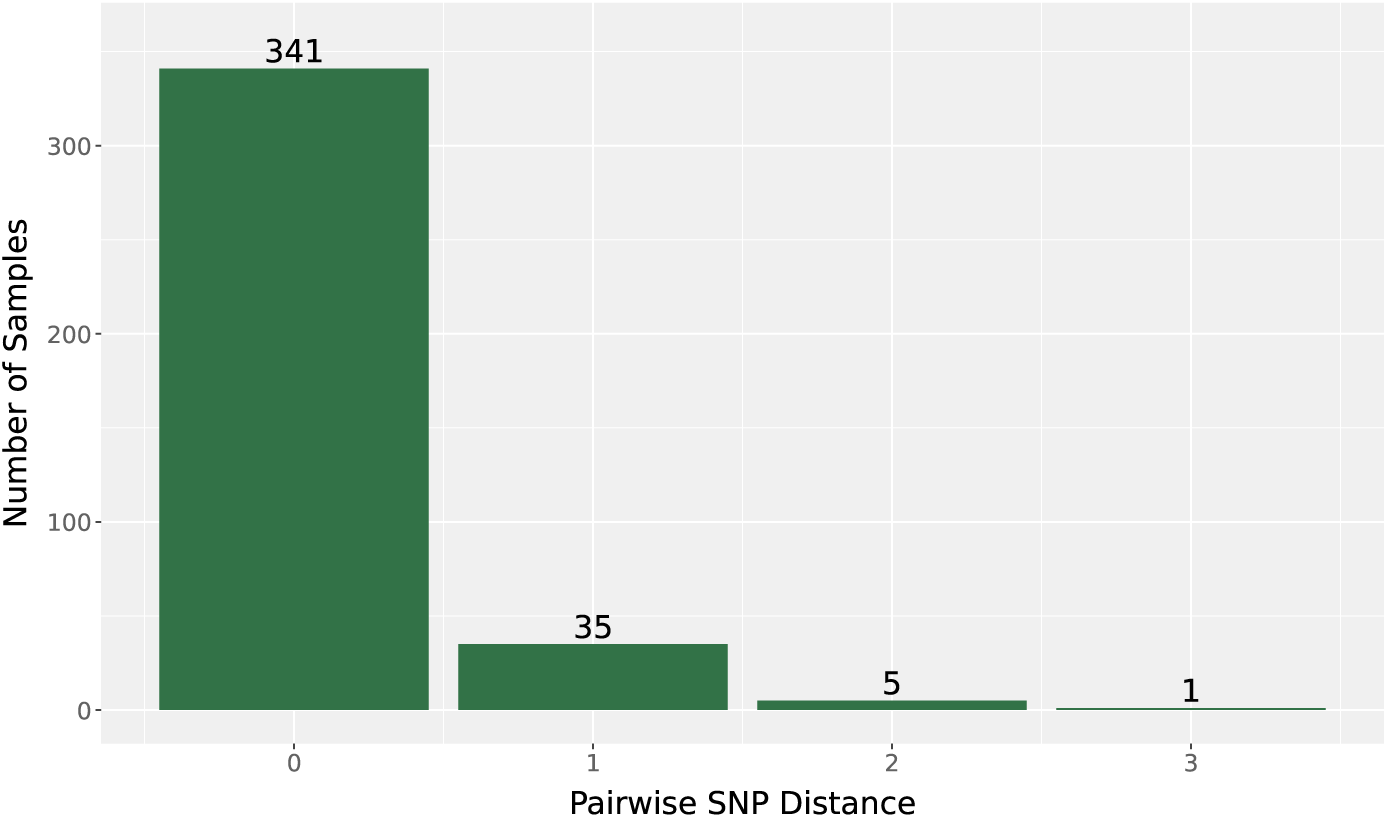
SNP distance between ONT and Illumina assemblies on the same sample.

### Both sequencing technologies identify nearly identical epidemiological clusters

Potential transmission clusters were generated using either ONT or Illumina sequencing by joining samples that were 12 or fewer SNPs apart [25]. A total of 43 clusters were identified in this way and contained 172/382 (45.0 %) samples with the majority of clusters being identical in both platforms (Fig. 4). There were 29 clusters containing just a pair of samples, of which one was exclusive to Illumina and one to ONT. All 12 medium-sized clusters (three to five samples) were identical in both platforms. The second largest cluster (16 samples) is identical, apart from sample 212 which is only connected to the cluster by ONT. The largest cluster (54 samples) comprises highly-resistant lineage 2.2.1 samples from South Africa. It consists of two highly related sub-clusters connected via sample 219. The lack of links between the two sub-clusters suggests that this central sample may have null calls (in both platforms) at sites which distinguish the two sub-clusters. The near-completeness of the sub-clusters means that all samples are included by both sequencing technologies.

**Figure 4:**
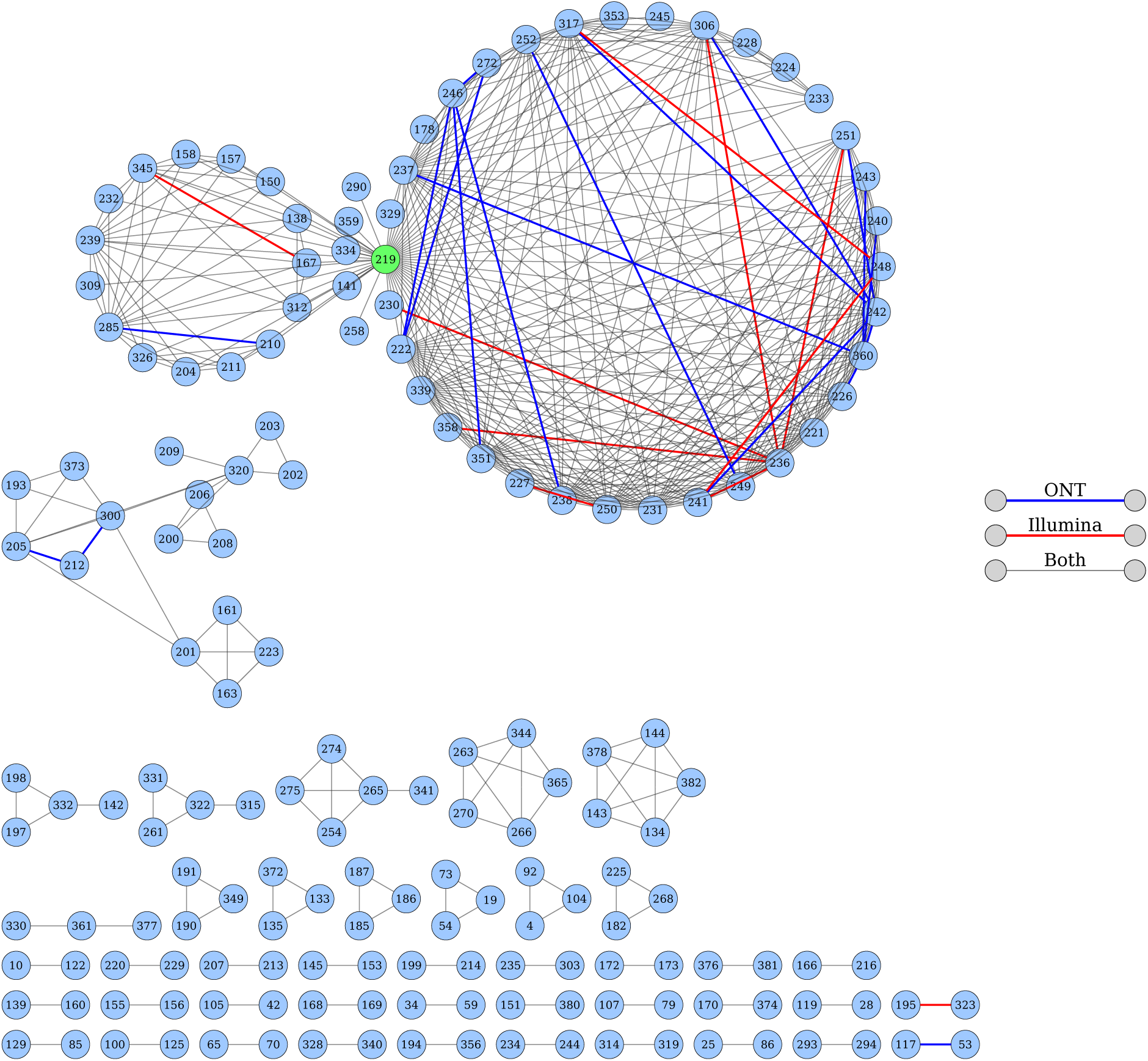
Transmission network produced at the 12 SNP threshold show broad agreement between platforms. Edges indicate which platforms include that connection. A black edge indicates that the link was made when the samples were sequenced with either technology whilst a red or blue edge indicates the link was only made when the samples were sequenced using either Illumina or ONT platforms, respectively. The “central sample” of the largest cluster is coloured green.

Comparing networks is crude as differences only arise when individual distances cross the 12 SNP threshold; the individual SNP distances could vary and we would be unaware. We therefore considered, for each pair of samples, the range of the four SNP distances that arise from the four permutations of the two sequencing platforms (Fig. S6 and S7). Fig. 5 displays the various SNP-distances for all the edges in Fig. 4.

**Figure 5:**
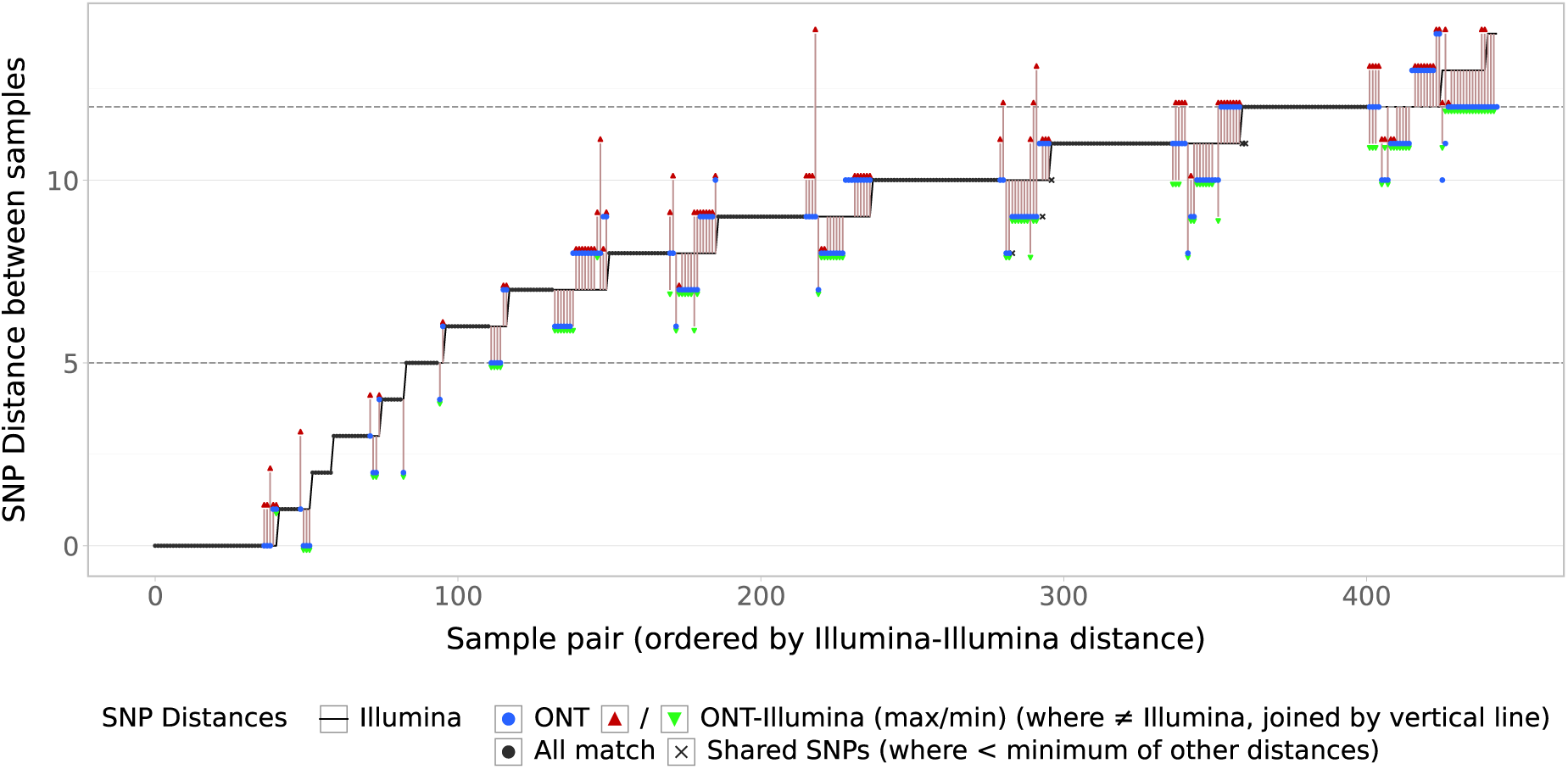
Variation in SNP-distance between technology. Pairs of samples (corresponding to the edges in Fig. 4) are ordered along the *x*-axis with the solid line indicating the Illumina SNP-distance. Blue dots show ONT distance, the red/green triangles show the max/min cross-technology distance (only plotted if different to Illumina) with the two points being joined by a vertical line. Where all distances are the same, just a black dot is drawn. The crosses indicate shared SNPs (the number of SNPs common to all four comparisons) and are only drawn if less than the minimum of the other distances.

We found ONT distances to be similar to Illumina and identical in the majority of cases. Notably there are no cases where the Illumina distance is five SNPs or fewer but the ONT or cross-platform distance is greater than 12 SNPs. The range of the four SNP distances was one or fewer in 400/444 (90.2%) edges with the maximum of five only occurring once (Fig. 6).

**Figure 6:**
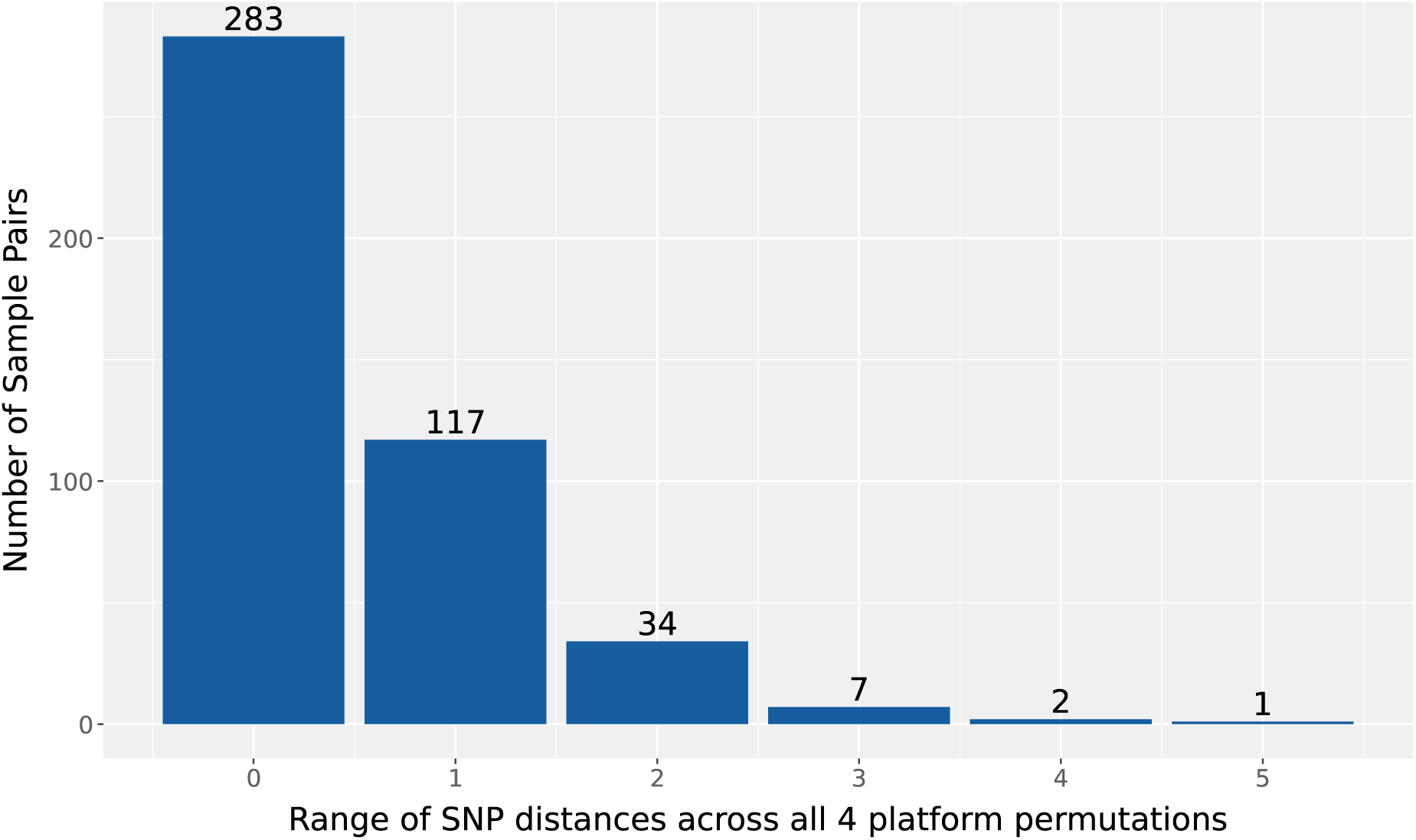
Range of SNP-distances between the four platform permutations for all sample pairs in the 12-SNP network. Each sample pair corresponds to an edge in Fig. 4.

## Discussion

This is the largest study to date investigating the ability of long-read sequencing (Oxford Nanopore Technologies) to resolve the key features of public health and clinical interest from clinical samples taken from patients with tuberculosis. The dataset of 425 isolates was collected in two countries (South Africa and Vietnam), included all the main *M. tuberculosis* lineages and was over sampled for antibiotic resistance. Strains yielded data on resistance prediction for 15 drugs and relatedness on 43 clusters among 382 strains. Using Illumina sequencing as the *de facto* reference standard, we found sufficient concordance in the detection of mixtures, the prediction of antibiotic resistance and membership of epidemiological clusters to conclude that long-read sequencing (specifically ONT flowcells R10.4.1 with V14 chemistry basecalled using dorado) has reached the point of maturity whereby ONT is a suitable alternative to Illumina sequencing. Our results provide evidence not only for the adoption of ONT by clinical or public health services but, equally importantly, that datasets made up of samples sequenced either by Illumina or ONT can be analysed in the aggregate to e.g. detect putative outbreaks of tuberculosis. This is a key advantage for public health TB genomics analyses at a country or international level. It also broadens the options for users who already have either sequencing technology or are seeking to invest in sequencers and are uncertain of comparability. The concordant performance of the two sequencing platforms we observe likely arises from the recent substantial improvements in ONT flow cells, chemistry, and base calling and variant calling algorithms.

Due to the likelihood of multiple infections and latency, tuberculosis infections are often mixtures and, since clinical samples are usually cultured in liquid media, genetic sequencing using Illumina platforms has been shown to detect both mixtures [26] and subpopulations containing resistance-associated variants [27, 28]. The latter have been shown to confer antibiotic resistance and our data demonstrate that ONT identifies fewer of the minority variants. This is due to a combination of inherent lower read depth, our higher minimum read threshold for ONT (as the error rate remains higher than Illumina) and the reluctance of Clair3 to call minor variants. These limitations can be mitigated by increasing the read depth of ONT sequencing so it is on a par with Illumina (e.g. 50 fold depth), improved accuracy and the continued inevitable improvements in machine learning algorithms like Clair3 for variant calling.

The Clinical and Laboratory Standards Institute (CLSI) state that a proposed DST method should, compared to a reference method, have a very major error rate (VME, false negative rate) of ≤ 1.5 % and a major error rate (ME, false positive rate) ≤ 3 % [23]. Using Illumina sequencing as the reference method, and considering all 15 drugs together, the VME rate is 1.0% (0.6-1.5%) and ME rate is 1.7% (1.3-2.2%) with 6.9% (6.3-7.5%) of results not having a definite resistant or susceptible classification being returned and have therefore been excluded from the VME/ME calculation. Individually, all 15 antibiotics had VME rates ≤3% with 11 being ≤1.5%. Of those 11, six had 95% confidence intervals below 5% and the other five had no very major errors but too few resistant samples to meet this bound. ME rates were ≤1.5% for all drugs with confidence intervals below 5% except delamanid, owing to 60 discrepant results due to a *fbiC* tandem-repeat deletion triggering an inappropriate rules-based resistance call in ONT.

Genetic-based prediction of drug susceptibilities relies on catalogues of genetic variants and their associated effects on specific antibiotics; the current gold standard is the second edition of the WHO catalogue of mutations in *M. tuberculosis* (WHOv2) [22] which associates specific genetic variants with the results of phenotypic drug susceptibility tests. The dataset used to derive this catalogue entirely used samples sequenced using Illumina and consequently the performance of a large dataset sequenced with ONT is expected to be similar to those se-quenced via Illumina. Such catalogues are, of course, incomplete and have unfortunate but inevitable prediction gaps for new drugs where resistance is still evolving, notably bedaquline, pretonamid and linezolid – members of the new BPaL(M) regimen for treating multi-drug resistant tuberculosis [29]. If we focus on bedaquline, the WHO report a VME rate of 40.4% [22] and even building a specific catalogue that is more permissive and concentrates on classifying minor variants identified by Illumina sequencing only reduces this to 19.2% for the 95.1% of samples that reached a definite classification [30]. The VME rate will no doubt reduce as more resistant isolates are collected, improving our ability to make statistical associations, but the ability of ONT to better detect large structural changes, or at least rule them out, will help [4]. A large international consortium modelled on CRyPTIC is being initiated to study how resistance to the BPaL(M) regime develops which, along with other studies, provide valuable data for follow-up studies.

To investigate if ONT sequencing can determine if two samples are epidemiologically related, we first removed any samples with evidence of being mixtures, leaving 382 samples for analysis. Crucially, a mask (264,525 loci, 6% of the H37Rv genome) was developed to exclude genome regions with frequent systematic differences between the two sequencing platforms due to being repetitive or of low complexity. This yielded a mean SNP distance between replicates sequenced with both platforms for the same sample of 0.13 with 376/382 (98.4, CI:96.6-99.4%) being ≤1 SNP apart. This is a major improvement on the 0.75 result reported by Hall *et al* [5]. Consequently, only minor differences are found amongst the 43 putative outbreak clusters (representing 172 isolates) identified using a 12-SNP threshold, with 40/43 clusters being identical regardless of the mix of the two platforms used to produce them.

Our study has several shortcomings; notably it is retrospective and assumes that Illumina sequencing can be treated as a reference method. To meet current regulatory standards ONT sequencing needs to be validated prospectively in the field, including collecting and comparing to phenotypic drug susceptibility data as the reference standard. Furthermore, the relatively small number of samples sequenced via ONT platforms also constrained our statistical resolving power in places and also meant we could not independently validate the genomic mask we built.

Overall, however, the ability of ONT sequencing to resolve the relevant public health and clinical information from clinical *M. tuberculosis* samples is hugely encouraging and, whilst our conclusions are not transferable to other pathogens, should increase the confidence of those wondering about applying ONT sequencing to a wider range of pathogens.

## Contributors

DWC, TEAP and PWF conceptualised the study. CSB, AH, DWC, HW and ER planned and comissioned or executed sequencing. MC performed the comparison of sequencing results. HNH, PPT, DDAT, NLQ, NTTT, TMW and RS sourced all samples from Vietnam. SVO sourced all samples from S. Africa. MC, JW, HT, RDT, and PWF developed the bioinformatic pipeline used for analysing the results. MC, CSB, ER, DWC, TEAP, PWF and RDT wrote the manuscript. All authors reviewed and agreed to submission of the manuscript.

## Data sharing

Reads have been deposited in the ENA (PRJEB106186) and will be released on acceptance and the project code added at proof.

## Supporting information

supp_mat_6_indel_variants

supp_mat_5_variants_on_536_res_sites

supp_mat_4_mutations

supp_mat_3_sample_pairwise_comparison

supp_mat_2_sample_filtering_counts

supp_mat_1_manifest_metadata

## Acknowledgements

The authors would like to acknowledge funding from the NIHR Health Protection Research Unit: Healthcare Associated Infections and Antimicrobial Resistance at University of Oxford (NIHR200915), a partnership between the UK Health Security Agency (UKHSA) and the University of Oxford, the National Institute for Health and Care Research (NIHR) Biomedical Research Centre: Oxford and the Ellison Institute of Technology, Oxford Ltd. For the purpose of open access, the author has applied a CC BY public copyright licence to any Author Accepted Manuscript version arising from this submission. The findings and conclusions in this report are solely the responsibility of the authors and do not necessarily represent the official views of the NHS, the NIHR, UKHSA, the Department of Health and Social Care or the Ellison Institute of Technology, Oxford Ltd.

