## supplementary_material for "Long-read sequencing of *Mycobacterial tuberculosis* is comparable to short-read sequencing for antimicrobial resistance prediction and epidemiological studies"

### 1 Description of supplementary materials

The following supplementary tables have been provided:

1. The mycobacterial references used in the competitive mapping step.
2. Sample filtering counts, split by flow cell.
3. Summary table of all samples with main pairwise comparison results.
4. All mutations found in resistance associated genes.
5. Comparison of variants on the 536 high-confidence resistance associated loci.
6. All indel variants across the genome.

Note: In table headings Illumina is referred to by the suffix "_1" or "A", and ONT by "_2" or "B"

Table S1: Description of flowcell runs for ONT.

| Flowcell | Samples | Dorado version | Sequencer |
|---|---|---|---|
| South Africa 1-13 | 309 | 4.3.0 SUP | GridION |
| Vietnam A,B (commercial supplier) | 192[1] | 5.0.0 SUP | PromethION |
| Vietnam 1 | 20 | 4.3.0 SUP | GridION |
| Vietnam 2,3 | 48 | 5.0.0 SUP | GridION |
| South Africa 14 / Vietnam 4 | 24 | 4.3.0 SUP | GridION |

[1] 81 of these were repeated in Vietnam 1-4

#### 2 Sample filtering

We collected 508 samples from South Africa and Vietnam with existing Illumina sequencing and successfully sequenced 502 (six failed PCR amplification). Three out of the 19 flow cells had substantially lower yields (<100 Mbp per sample) and we therefore repeated the 81 samples most affected. We filtered out samples with <10X read depth and with <90% resolved genome coverage (proportion of the genome which is not called null, noting that deleted sites are not considered as null for this purpose), resulting in 425 samples for comparison (Fig. S1, S2).

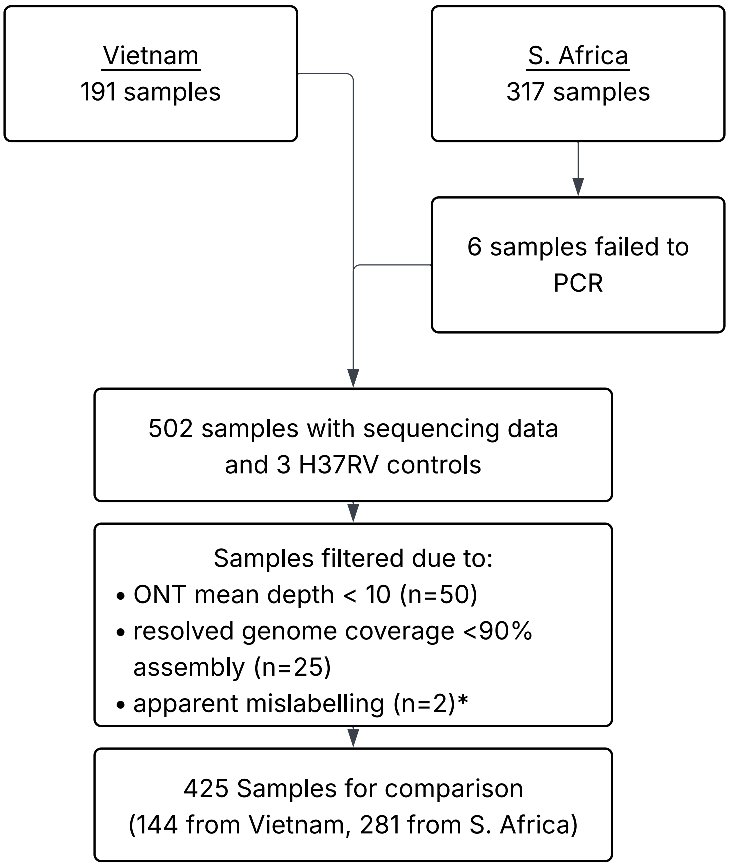

Figure S1: Sample collection and filtering.

*Samples differed greatly between platforms in lineage, AMR, and relatedness - far more than any other samples.

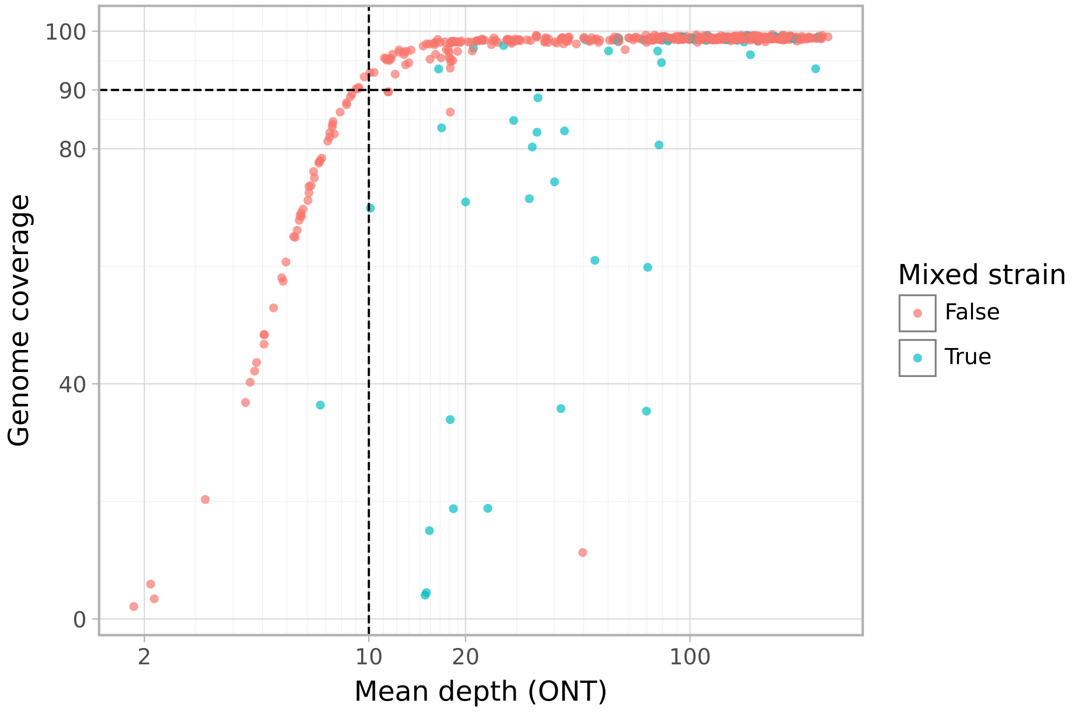

Figure S2: ONT mean depth (log scale) vs resolved genome coverage. Samples are coloured based on being mixed strain (see supplement §6). The 10X depth and 90% coverage filters are marked as dashed lines.

### 3 Sequencing quality statistics

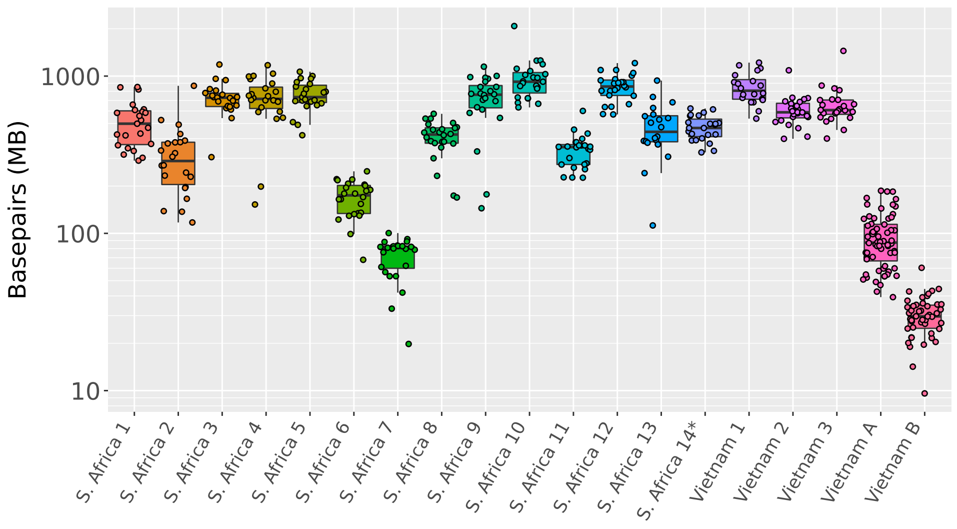

(a) ONT sequencing quantity

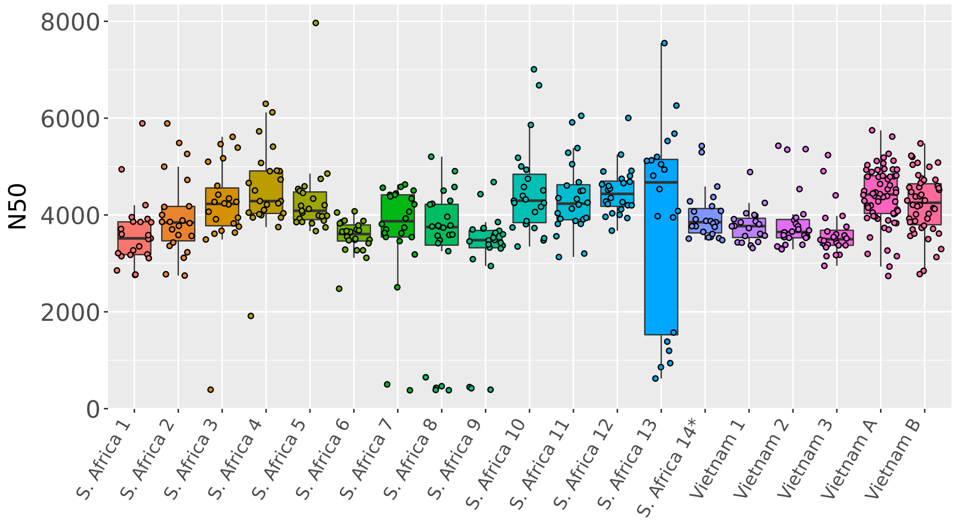

(b) N50 distribution

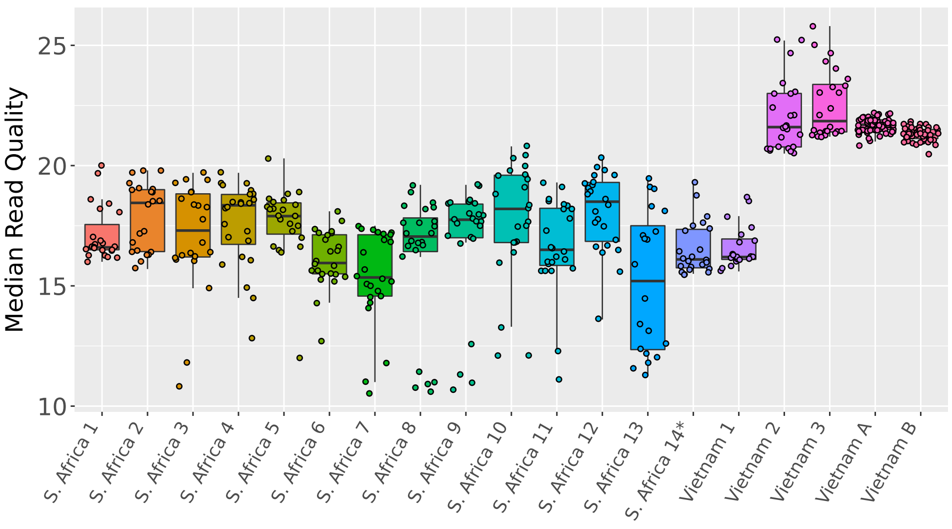

(c) Read quality distribution

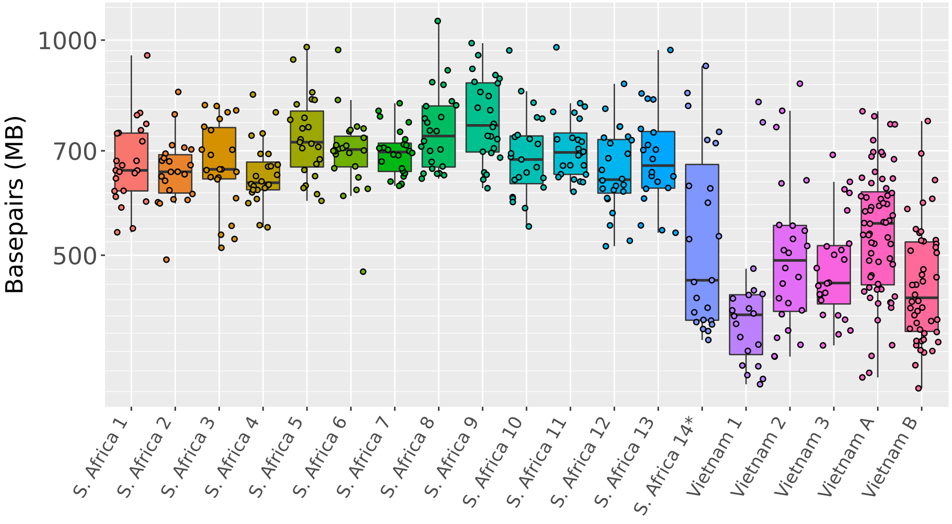

(d) Illumina sequencing quantity

Figure S3: Summary of sequencing statistics by ONT flowcell.

*The S. Africa 14 flowcell was also used for the Vietnam 4 batch. Vietnam 2,3,A,B used dorado SUP 5.0.0, the rest used SUP 4.3.0

### 4 Species, Lineage, and subpopulation

Species was determined by mapping reads against a collection of mycobacterial reference genomes, which also enabled filtering for reads mapping against H37Rv. We arbitrarily applied a threshold of 40% genome coverage to signify "strong support". To detect potential minor subpopulations we used a threshold of 5% genome coverage and required that the genome coverage was reasonable given the mean read depth. This was done to limit the effect of read-misassignment from the major species. To do this we estimated the expected coverage using a Poisson distribution on the mean depth, and then required the actual genome coverage to be at least 50% of the expected coverage.

$$\text{expected coverage} = 100 \times (1 - \exp(-\text{mean depth}))$$

Mykrobe was used to determine TB lineage. Calls with the filter `LOW PERCENT COVERAGE` were excluded due to being exclusively lineage 4.10 in low depth samples.

Of the 426 samples which passed quality filtering, 18 were found to have mixed lineages, eight were found to have evidence of a non-tuberculosis mycobacteria (NTM) subpopulation, and one sample had both mixed lineage and a NTM subpopulation. Of the mixed lineage samples:

- 16 samples had a mixed lineage according to Illumina with ONT only reporting one of the lineages.
- one sample was lineage 2.2.6/4.4.1.1 in Illumina but only lineage 4 in ONT, most likely due to low read depth in ONT.
- two samples were lineage 2.2/2.1 in ONT but only lineage 2.2 in Illumina. Notable, in both samples, the read support for the 2.1 call was very low with a reported "percent_coverage" below 25%.

Of the samples with NTM subpopulations seven had *M.avium*, one had *M.novum* and one had *M.marseillense*. Read mapping in ONT gave strong support for all nine, while Illumina gave strong support for 5 and weak support for 4. Mykrobe also reports species, but was less sensitive than read mapping. It found 6/9 subpopulations in ONT, and only 2/9 in Illumina.

### 5 Variant Calling

Variant calling was performed with Clockwork for Illumina and Clair3 for ONT. In addition BCFTools was used to report read depths at sites not included in the Clair3 VCF. The minimum number of reads required to support a variant call was three in Illumina and five in ONT. A variant was deemed "major" if it had $\geq$90% of the read depth at that site, but would still be considered as a "minor" variant for resistance prediction provided it had a ≥5% of the read depth. If there was no major call at a site then it is set to null (N) in the fasta file. Note that no SNPs or minor alleles were called based on the BCFTools results and was only used for calling nulls.

Filters were also applied in addition to the read depth requirements already mentioned. Clair3 VCF rows required quality scores ≥2. BCFTools required mapping quality scores ≥30. Clockwork filtered rows with a depth three standard deviations above the mean depth as well as sites with a genotype confidence score below 0.5. Filtered rows resulted in null (N) calls in the fasta file and variants from such rows were not used in resistance prediction.

We noted a greater proportion of indels were classified as minor in ONT than in Illumina, likely as a result of noisier reads reducing the proportion of them matching the indel allele.

### 6 Detecting mixed strains

To detect mixed strains (even in cases of identical lineage) we counted the number of mixed sites within a sample. These are sites with support for two alleles (one of which may be the reference allele). Note that we did not consider indels in this analysis. Support in this context means at least 3 reads in Illumina or at least 5 reads in ONT, as well as being at least 5% of the read depth at that site. Mixed sites within 12 basepairs of other mixed sites were excluded (in the same way as when calculating SNP distances).

Samples labelled mixed lineage by Illumina show an elevated but varied number of mixed sites by both platforms (Fig. S4) although ONT reports fewer calls in some samples which may be linked to low depth. The two samples labelled mixed lineage in ONT only show no evidence of being mixed in Illumina (with only 6 and 11 mixed sites) despite evidence in ONT (124 and 193 mixed sites).

We decided to use 124 mixed sites (the minimum number of mixed sites within the mixed-lineage samples) as the cut-off for considering a sample to have evidence of being a mixed strain. Using this there are 43 samples with some evidence of being mixed strains: 21 in both (including the 17 Illumina mixed lineage samples), four in Illumina only, and 18 in ONT only (including both ONT mixed lineage samples). Notably all ONT-only mixed samples came from the S. Africa collection making it plausible that it's not just a bioinformatic issue with detecting minor SNPs.

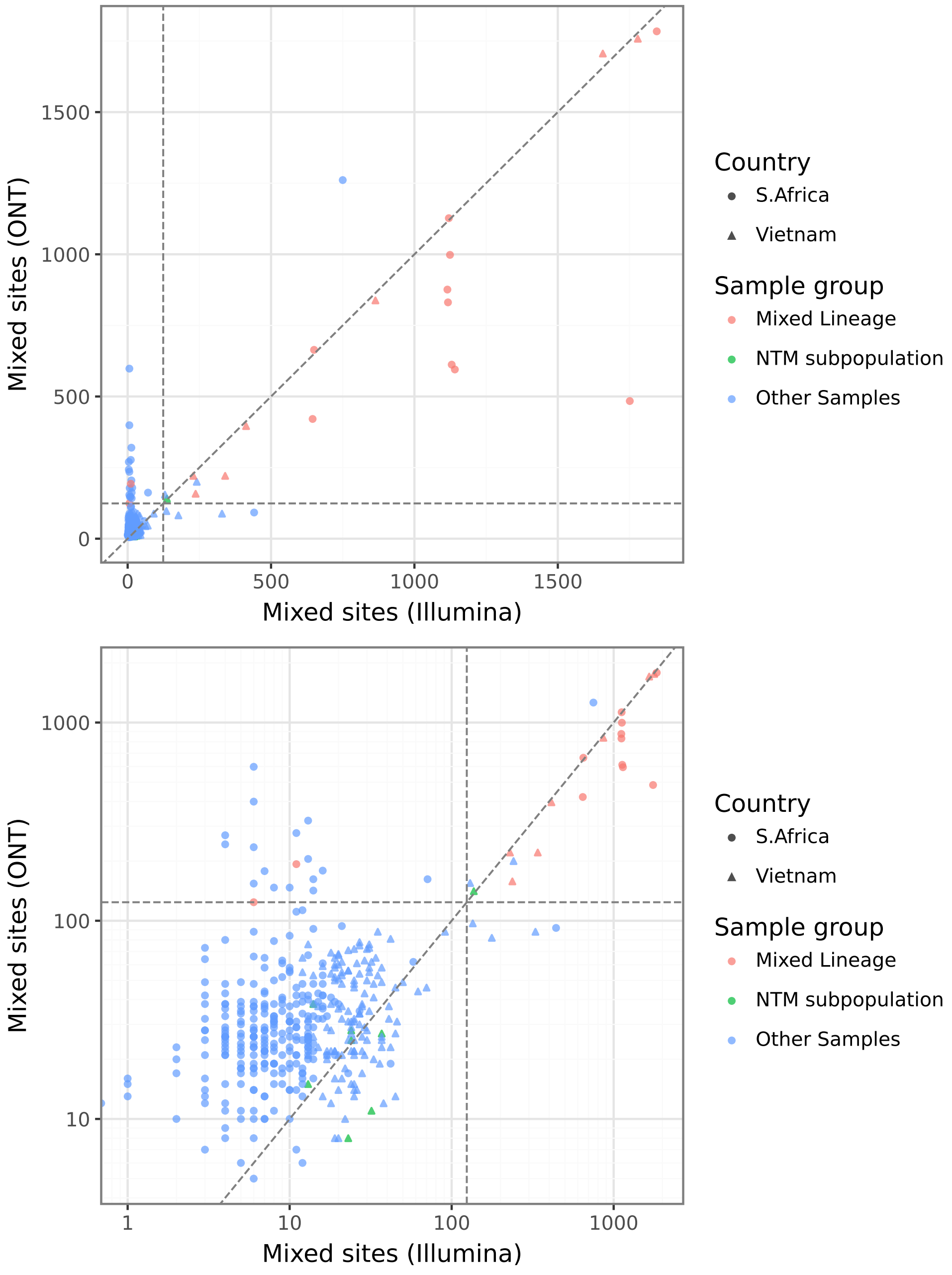

Figure S4: Counts of mixed sites between platforms with linear axis (a) and log axis (b). The 124 mixed sites cut-off is marked with a dashed line.

### 7 Antibiotic resistance detection

Note that there was a single mutation which was represented as two SNPs in Illumina and as a 1bp deletion and insertion in ONT. We have decided to count this as two matching SNPs as they do not represent true platform differences.

Table S2: Confusion matrix of variants found on the 536 high-confidence resistance loci

| | ONT | | | | | | |
|---|---|---|---|---|---|---|---|
| Illumina | ref | null | SNP | minor SNP | del | minor del | total |
| ref | 225,874 | 21 | 0 | 1 | 0 | 13 | 225,909 |
| null | 0 | 6 | 0 | 0 | 0 | 0 | 6 |
| SNP | 0 | 0 | 1,320 | 18 | 0 | 0 | 1,338 |
| minor SNP | 26 | 0 | 15 | 102 | 0 | 0 | 143 |
| del | 0 | 6 | 0 | 0 | 279 | 75 | 360 |
| minor del | 0 | 0 | 0 | 0 | 0 | 41 | 41 |
| 2 minor SNPs[1] | 3 | 0 | 0 | 0 | 0 | 0 | 3 |
| total | 2259,03 | 33 | 1,335 | 121 | 279 | 129 | 227,800 |

[1] these positions had two different alternate alleles.

Table S3: Confusion matrix of resistance-associated mutations (or those with unknown classifications) found within resistance-associated gene

| | ONT | | | | | | | |
|---|---|---|---|---|---|---|---|---|
| Illumina | amino acid mutation | minor amino acid mutation | indel | minor indel | null | complex | absent | total |
| amino acid mutation | 2,168 | 29 | 0 | 0 | 0 | 0 | 0 | 2,197 |
| minor amino acid mutation | 26 | 103 | 0 | 0 | 0 | 0 | 25 | 154 |
| indel | 0 | 0 | 191 | 49 | 0 | 0 | 2 | 242 |
| minor indel | 0 | 0 | 0 | 50 | 0 | 0 | 27 | 77 |
| null | 0 | 0 | 0 | 0 | 0 | 0 | 0 | 0 |
| complex | 0 | 0 | 0 | 0 | 0 | 2[1] | 0 | 2 |
| absent | 0 | 1 | 0 | 63 | 0 | 0 | 0 | 64 |
| total | 2,194 | 133 | 191 | 162 | 0 | 2 | 54 | 2,736 |

[1] In both instances ONT and Illumina shared a single SNP/minor SNP but Illumina had an additional minor SNP within the same codon leading to different amino acid mutations. In one case this caused ONT to predict resistant whilst Illumina called susceptible. The other case both codon mutations triggered a resistant call.

##### 7.1 Deletions in *fbiC*

The end of the *fbiC* gene is a tandem repeat with a 62bp section of DNA which occurs twice in full and finally a section of 43bp. In 23 of our samples, the ONT assemblies had one of these repeats deleted. Since each 62 bp section contains the *fbiC* stop codon the deletion of just one of the repeats does not therefore alter the resulting primary protein sequence.

##### 7.2 2 kbp deletion in *katG*

One of the samples had a 2 kbp deletion in *katG* which was reported by Illumina, but not by ONT. However, the deletion was clear in the ONT read pileup, but clair3 did not call the variant. Instead all the sites within the deleted range had low depth null calls.

##### 7.3 Indels across the genome

We investigated the difference in indel calling across the genome between Illumina and ONT. To do so all indels were first broken down into simple insertions or deletions. Any indel contained (or partially contained) in our mask was ignored. Each indel was considered 'filtered' if the VCF row had a filter flag (except for `MIN_FRS`, `OVERLAP_BETTER_VARIANT`, `MIN_GCP` which are often present in mixes).

For each indel called by one platform we checked if it was present in the other platform. If not then we checked for a 'clash' with another indel, 'null' calls in the other platform, or 'ref' otherwise. This matrix is presented in tables S4, S5.

Illumina reports 22231 major insertions of which ONT finds evidence for 21721/22231 (97.7%) meaning that it was present in ONT in some form (major, minor or filtered). Whereas Illumina only finds evidence for 18182/20011 (90.9%) ONT major insertions with 1637/1829 (89.5%) of the misses being due to null calls.

Deletions are similar with ONT finding evidence for 23239/23824 (97.5%) major Illumina deletions with 513/585 (87.7%) of misses being due to 'clashes' with other indels in ONT. Whereas Illumina finds evidence for 19685/25402 (77.5%) major ONT deletions with 5375/5717 (94.0%) misses being due to null calls. The null calls in this case may actually indicate the presence of the deletion in Illumina despite it not being picked up as a variant in the VCF.

From comparison with Illumina it is clear that ONT rarely missed indels which Illumina reports. The additional ONT indels may be spurious but note that many differences are present across many samples rather than being random errors.

| | ONT | | | | | | |
|---|---|---|---|---|---|---|---|
| Illumina | major | minor | filtered | ref | null | clash | total |
| major | 18,125 | 3,483 | 113 | 310 | 89 | 111 | 22,231 |
| minor | 44 | 764 | 10 | 336 | 155 | 22 | 1,331 |
| filtered | 13 | 0 | 0 | 38 | 5 | 12 | 68 |
| ref | 157 | 4,250 | 343 | 0 | 0 | 0 | 4,750 |
| null | 1,637 | 147 | 1,303 | 0 | 0 | 0 | 3,087 |
| clash | 35 | 85 | 48 | 0 | 0 | 0 | 168 |
| total | 20,011 | 8,729 | 1,817 | 684 | 249 | 145 | 31,635 |

Table S4: Differences in insertion variant calling

| | ONT | | | | | | |
|---|---|---|---|---|---|---|---|
| Illumina | major | minor | filtered | ref | null | clash | total |
| major | 19,524 | 3,634 | 81 | 30 | 42 | 513 | 23,824 |
| minor | 107 | 691 | 8 | 1,067 | 157 | 59 | 2,089 |
| filtered | 54 | 44 | 0 | 67 | 5 | 90 | 260 |
| ref | 151 | 1,704 | 333 | 0 | 0 | 0 | 2,188 |
| null | 5,375 | 1,038 | 7,506 | 0 | 0 | 0 | 13,919 |
| clash | 191 | 201 | 127 | 0 | 0 | 0 | 519 |
| total | 25,402 | 7,312 | 8,055 | 1164 | 204 | 662 | 42,799 |

Table S5: Differences in deletion variant calling

### 8 SNP distance

#### 8.1 Creation of a Mask for clustering

The SNP distance between two assemblies is the number of genome positions which have differing base calls. Positions where one or both assemblies have a null call are not included. A mask can be used when calculating SNP distances, by excluding certain sites from the calculation (note that no mask was used during the assembly step itself, only when calculating SNP distances). The aim of masking for relatedness is to exclude sites where SNPs arise as a result of incorrect mapping. The mask here is designed to cover low complexity and repetitive regions in the genome. We also incorporate work from Marin et al to mask regions with empirically poor Illumina mapping [1]. The mask is a compilation of the following methods:

1. Dust for low complexity.
2. mummer for tandem repeats.
3. NCBI annotation for direct repeats.
4. blastn for regions of self similarity.
5. Repeat regions from the NCBI GFF file (NC_000962.3).
6. Marin et al for empiric Illumina masking.
7. Small gaps in the mask ($\leq$100 bps) were added to the mask.

The resulting mask covered 264525 positions (6% of the genome). To assess the successfulness of this mask we looked at the SNP differences between platforms that were common across multiple samples (Fig. S5). The mask captured the majority of the discrepant sites but there were eight sites which had a SNP difference between platforms in at least ten samples. The eight sites were composed of a run of four sites, a pair of sites 2 bp apart, and two singletons. The 8 sites, along with the site in-between the pair, were added to the final version of the mask (bringing the total count up to 264534 positions).

Steps 1-4 and 7 can be easily applied to any genome fasta file. Steps 5 and 6 are specific to H37Rv.

A GitHub repository with the the code to reproduce the mask is here (`https://github.com/GlobalPathogenAnalysisService/masquerade`).

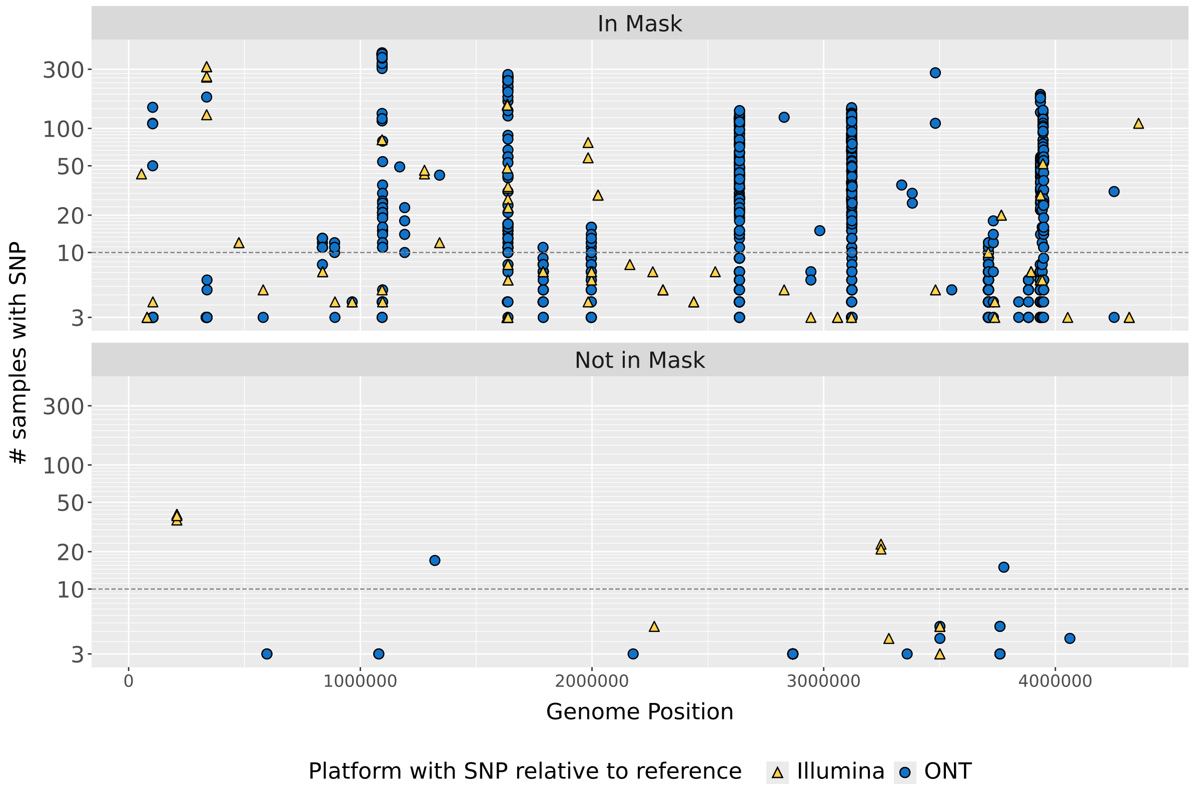

Figure S5: Manhattan plot showing distribution of SNPs across the genome between the Illumina and ONT assemblies. The y-axis indicates the number of samples that had a SNP at that site between the two platforms. The shape/colour indicates which platform had the SNP relative to the H37Rv reference. The top panel shows sites included within the initial version of the mask. The bottom panel shows sites not included in the mask. The line at $y = 10$ was used as the cut-off for selecting additional sites to be included in the final version of the mask. In the lower panel there are eight positions above the line although they are in four groups.

#### 8.2 Divergence between Illumina and ONT SNP-distance

We define the *min and max cross-technology-distance* as the lower and higher of the two SNP distances calculated between two samples when one is sequenced with Illumina and the other with ONT. We define *divergence* as the absolute difference between the Illumina distance and the ONT distance, and the *complex divergence* as the range of the four potential SNP distance metrics. Figure S6 and S7 show the 12-SNP transmission network (edges included if either ONT distance or Illumina distance $\leq 12$) coloured by divergence and simple-divergence. A sample's (complex) divergence is simply the mean of the samples edges in the transmission network.

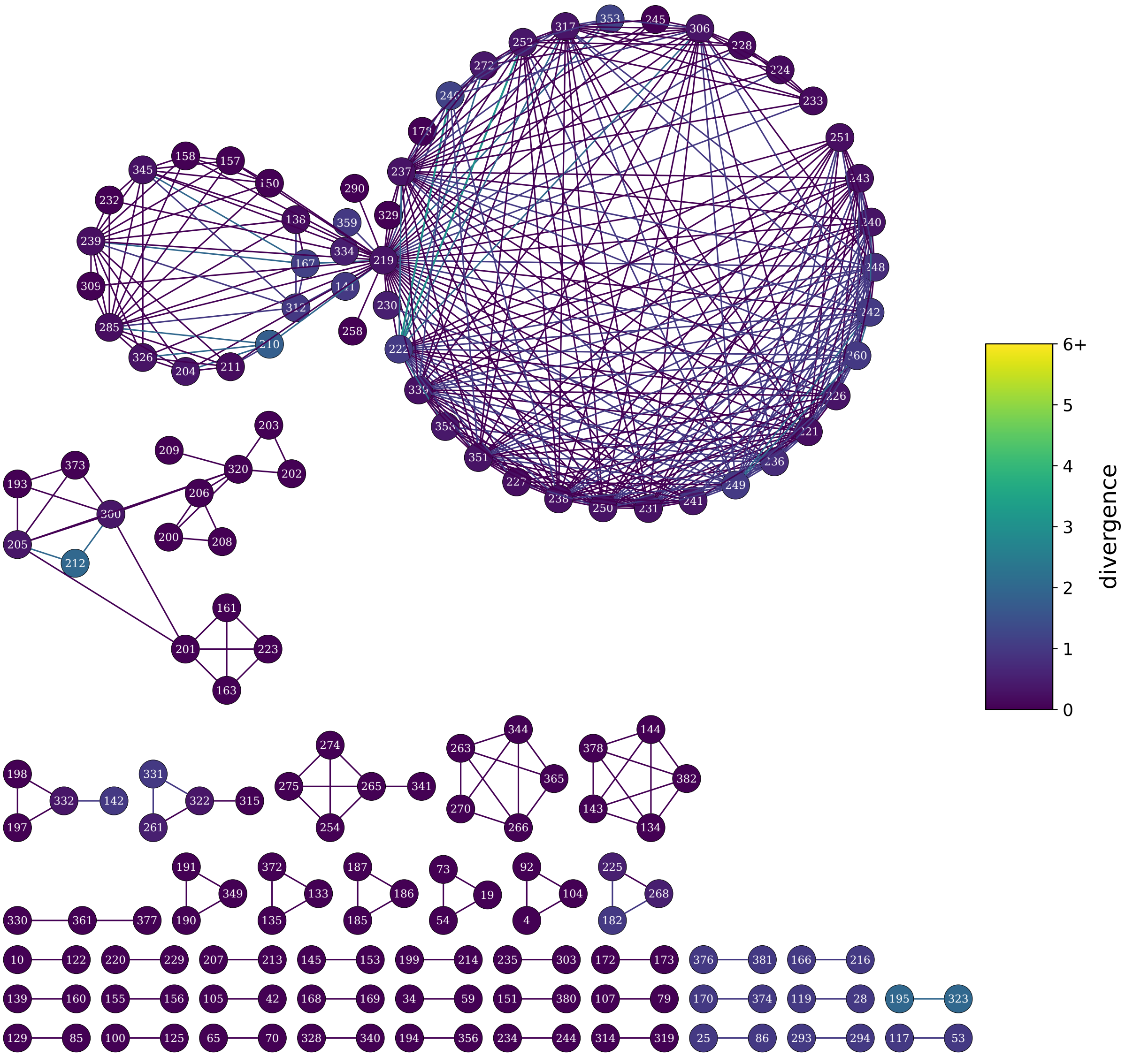

Figure S6: Transmission network produced at the 12 SNP threshold. Edges are coloured by divergence of SNP distances, while samples are coloured by the mean of their edges.

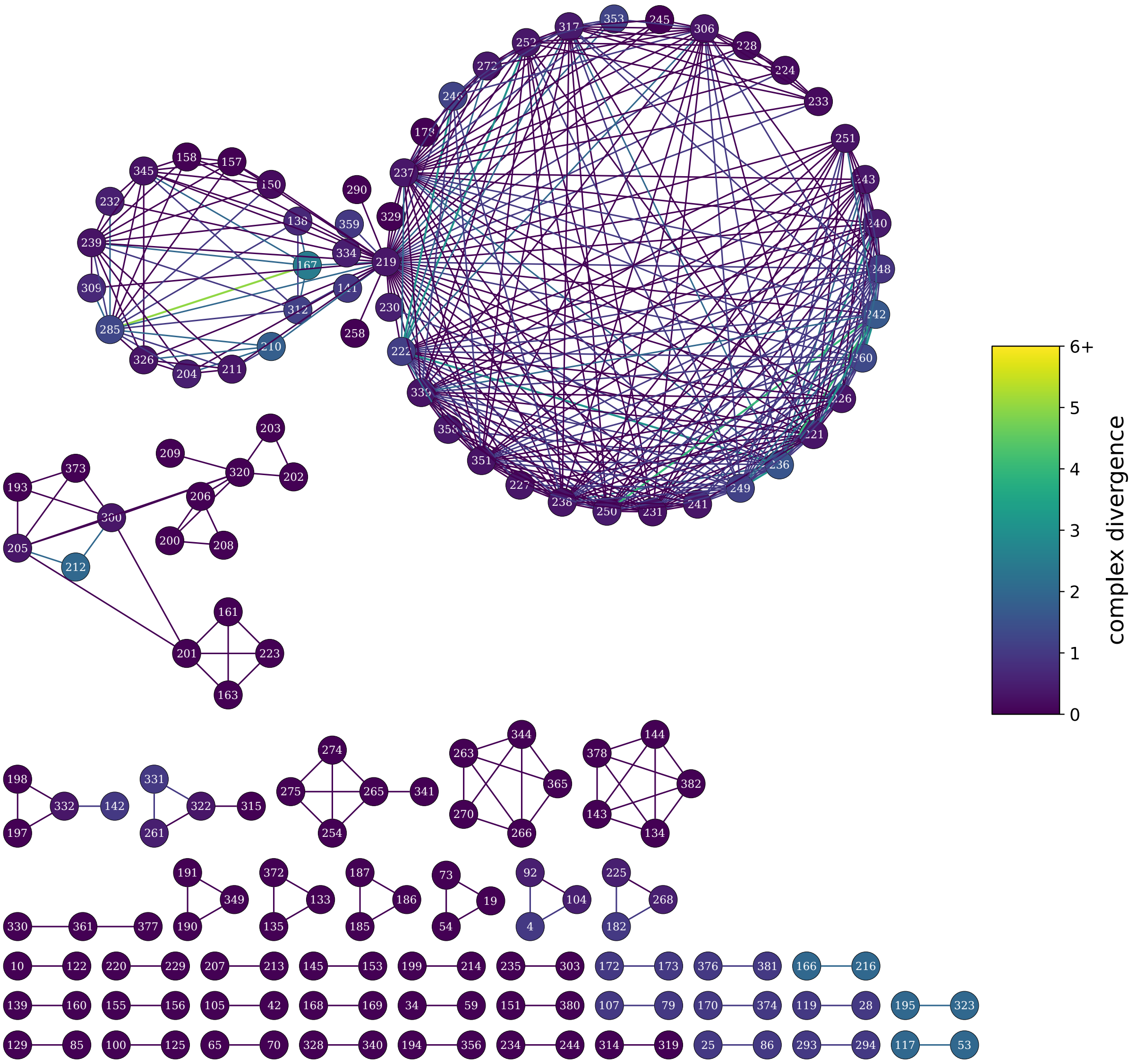

Figure S7: Transmission network produced at the 12 SNP threshold. Edges are coloured by complex-divergence of SNP distances, while samples are coloured by the mean of their edges.

#### 8.3 Impact of filtering nearby SNPs

A potential filter when calculating SNP distances is to exclude nearby SNPs, as a way of mitigating against spurious SNPs from mapping issues. Walker et al[2] use a distance of 12 base-pairs for this purpose, which is what we have used in this study. Specifically, for each assembled genome fasta we set any SNPs to N (null) if they are within 12 base-pairs of another SNPs. Here we are considering SNPs relative to the H37Rv reference. The result being that those sites are then masked for all comparisons with that assembly.

The filter has a clear benefit when comparing the distance between the Illumina and ONT assembly for the same sample (Fig S8). With the filter the largest discrepancy is 3 SNPs, but without this it goes up to 12 SNPs.

We wanted to know what effect the filter was having on clustering. Specifically if it caused a difference to which samples clustered using Illumina or ONT. We found that the filter did have an impact on which samples clustered at 12 SNPs; however, this was generally due to small changes near the 12 SNP threshold (see Fig S9,S10).

We also reran the clustering comparison between ONT and Illumina in the absence of the filter (see Fig S11, S12). We find that most pairs continue to have good agreement between platforms; however, we do find greater discordance both in pairs and in the larger clusters.

Even with high accuracy reads, there is the potential for a single poorly aligned section of the genome to throw off SNP-distance as a screening tool for transmission. As such requiring SNPs to be over 12 base-pairs from other SNPs still appears to be a useful filter.

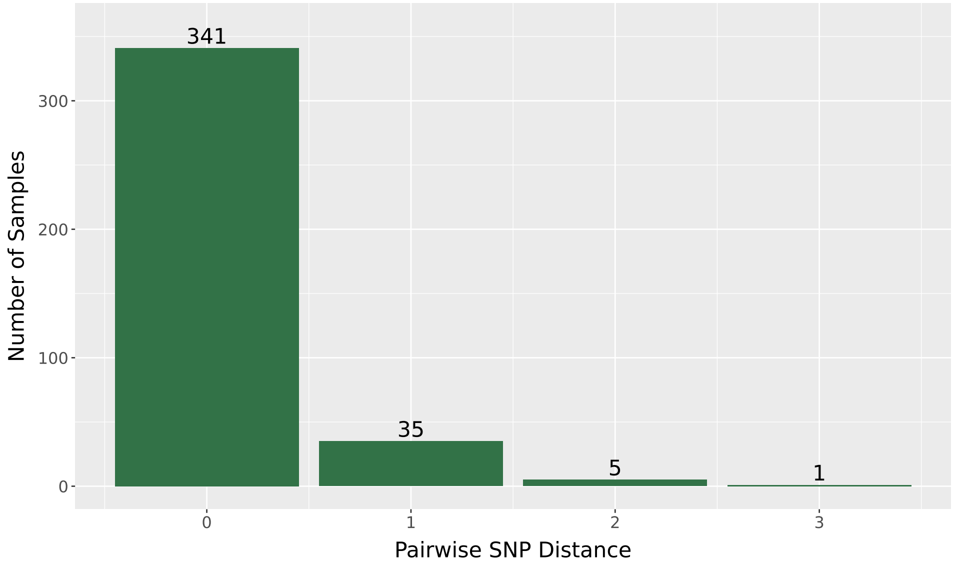

(a) SNP distances using 12 basepair filter

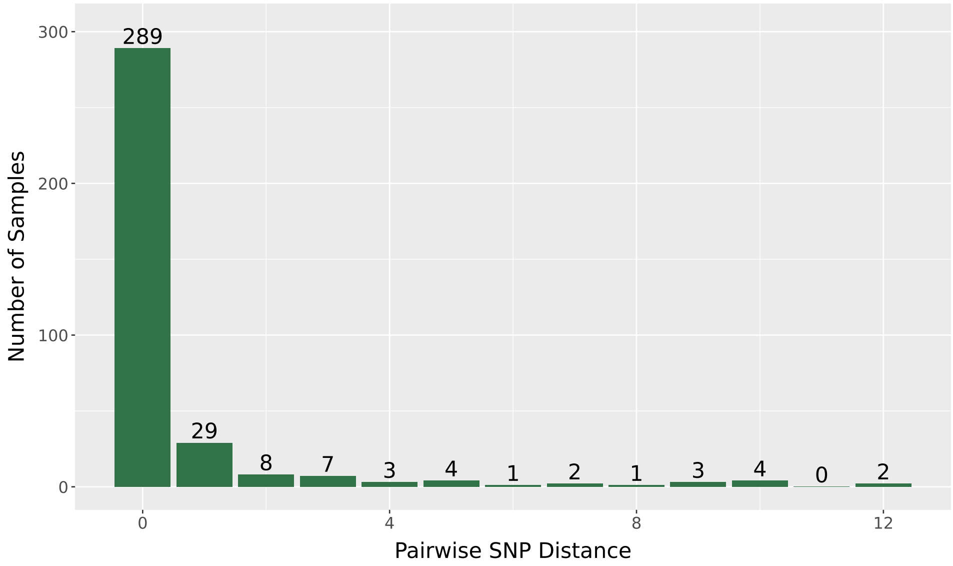

(b) SNP distances without 12 basepair filter. Note that an outlier (with distance 383) is not graphed.

Figure S8: Impact of SNP-distance filter on distance between Illumina and ONT on the same sample.

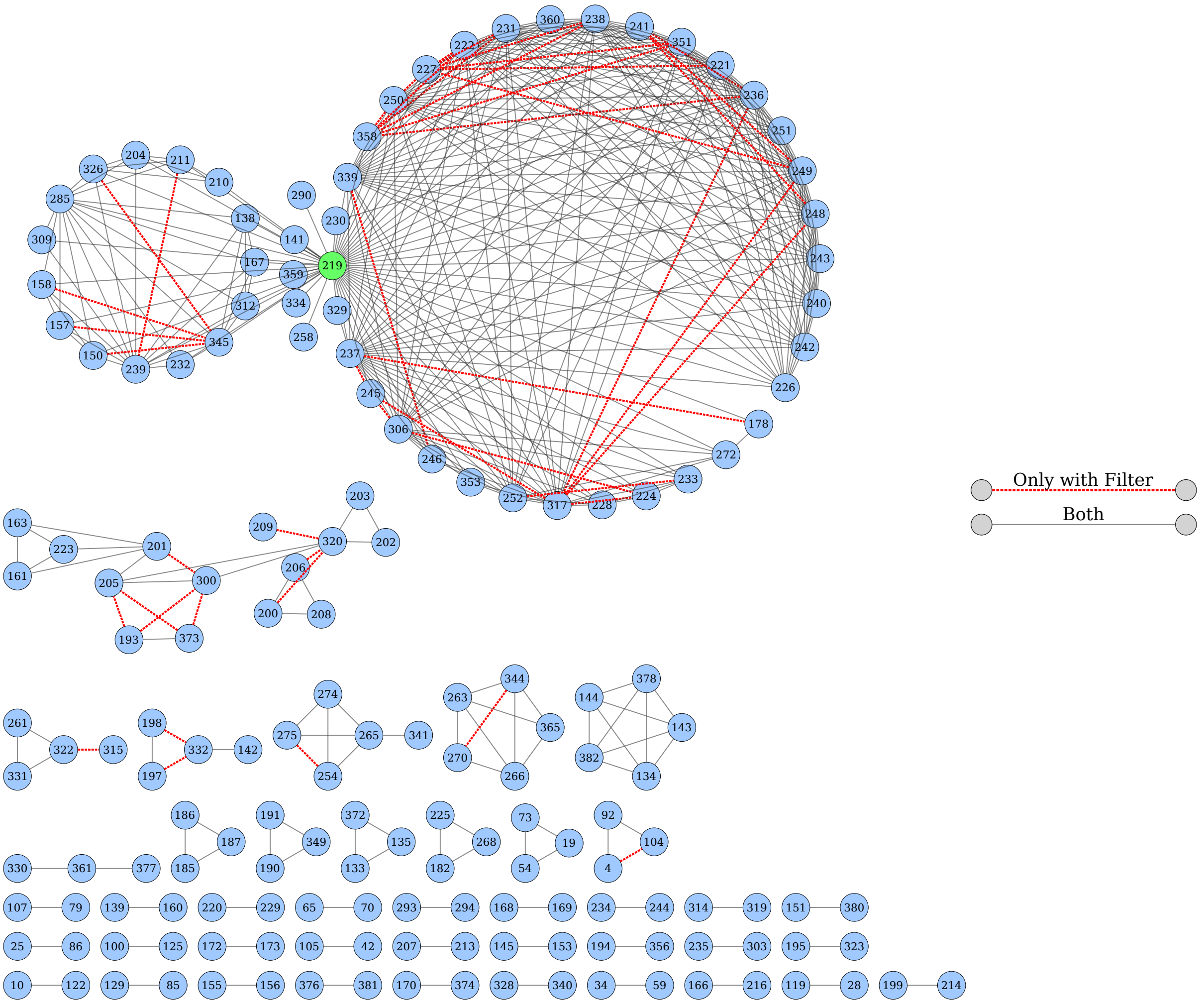

(a) Illumina relatedness with and without SNP-distance filter (using 12 SNP threshold).

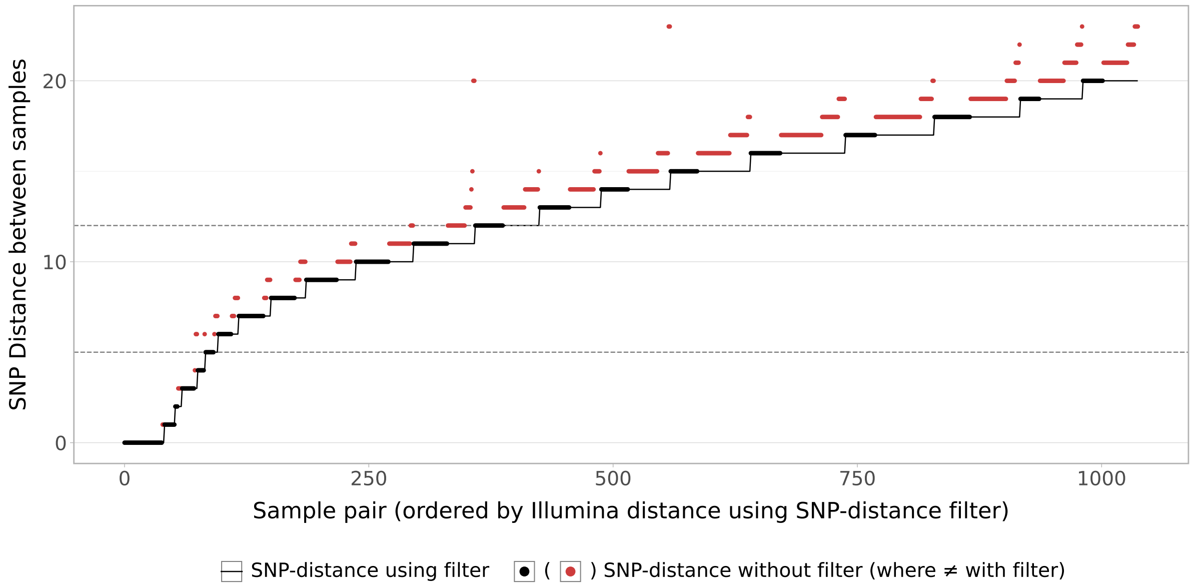

(b) Impact of SNP-distance filter on distance between pairs with Illumina.

Figure S9: Impact of SNP-distance filter on Illumina clustering

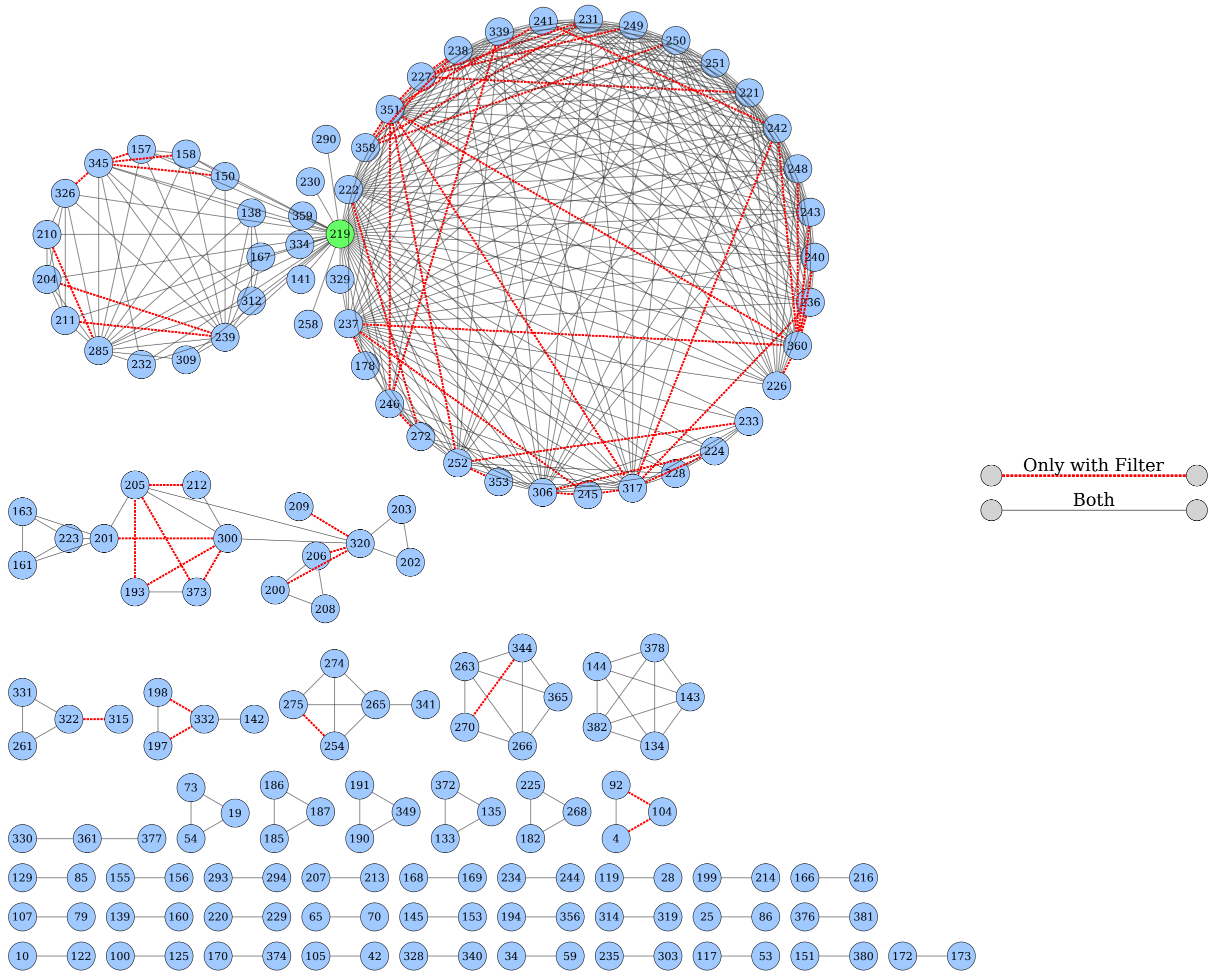

(a) ONT relatedness with and without SNP-distance filter (using 12 SNP threshold).

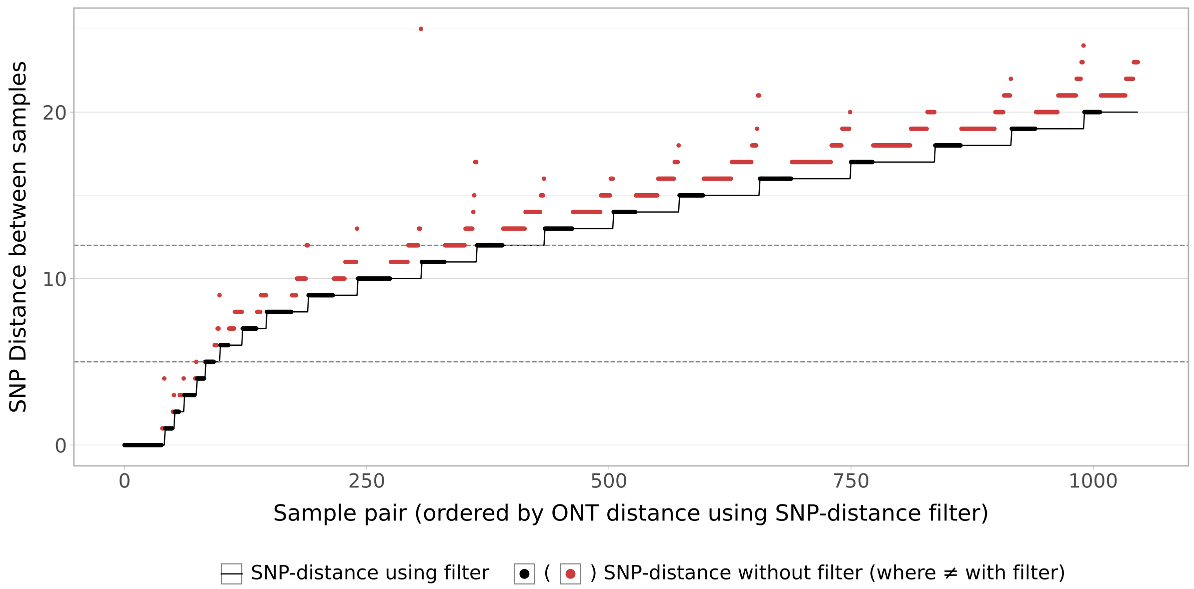

(b) Impact of SNP-distance filter on distance between pairs with ONT.

Figure S10: Impact of SNP-distance filter on ONT clustering

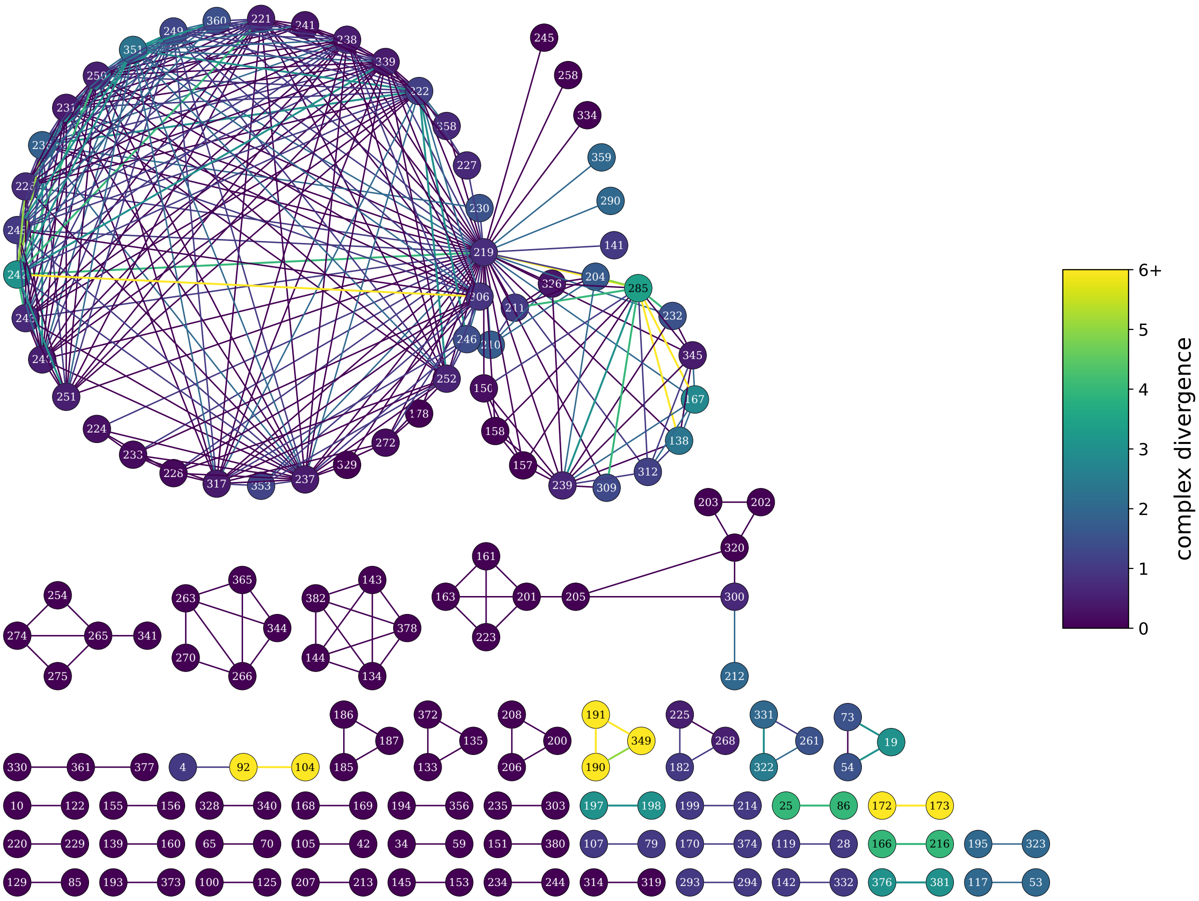

Figure S11: Transmission network produced at the 12 SNP threshold, without using SNP-distance filter. Edges are coloured by complex-divergence of SNP distances, while samples are coloured by the mean of their edges.

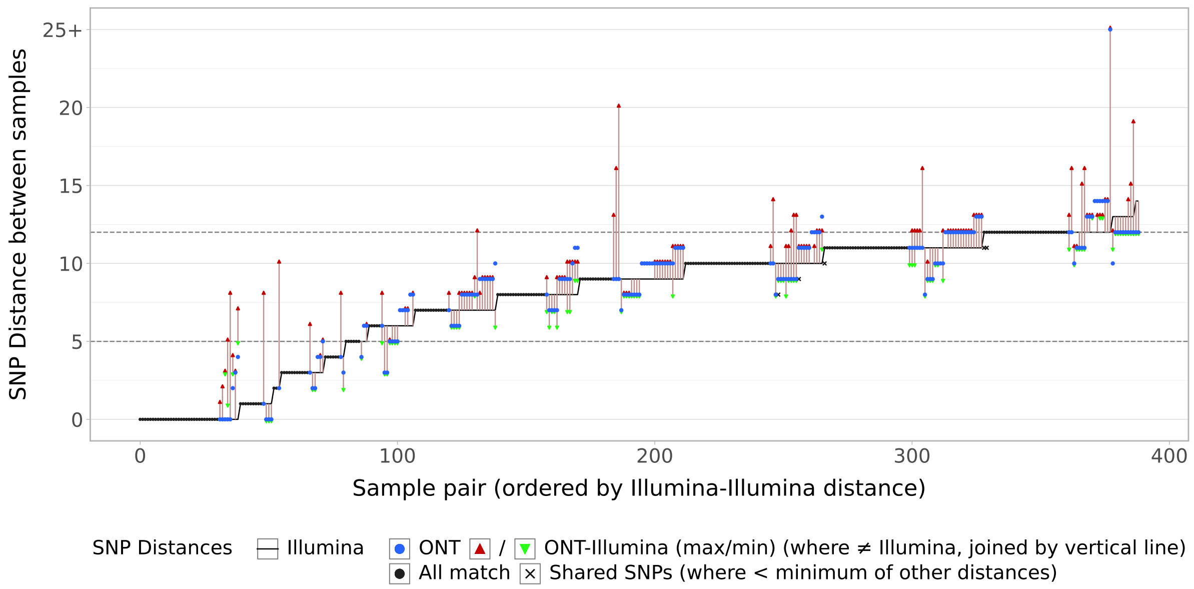

Figure S12: Variation in pairwise SNP-distance between platforms without using SNP-distance filter. Pairs of samples are ordered along the x-axis with the solid line indicating the Illumina SNP-distance. Blue dots show ONT distance, the red/green triangles show the max/min cross-technology (only plotted if different to Illumina) with the two points being joined by a vertical line. Where all distances are the same, just a black dot is drawn. The crosses indicate shared SNPs and are only drawn if less than the minimum of the other distances.
